# Cooperation between physiological defenses and immune resistance produces asymptomatic carriage of a lethal bacterial pathogen

**DOI:** 10.1101/2023.01.22.525099

**Authors:** Grischa Y. Chen, Natalia R. Thorup, Abigail J. Miller, Yao-Cheng Li, Janelle S. Ayres

**Affiliations:** Molecular and Systems Physiology Lab, The Salk Institute for Biological Studies, La Jolla, CA 92037; Gene Expression Laboratory, The Salk Institute for Biological Studies, La Jolla, CA 92037; NOMIS Center for Immunobiology and Microbial Pathogenesis, The Salk Institute for Biological Studies, La Jolla, CA 92037

## Abstract

Animals have evolved two defense strategies to survive infections. Antagonistic strategies include mechanisms of immune resistance that operate to sense and kill invading pathogens. Cooperative or physiological defenses mediate host adaptation to the infected state, limiting physiological damage and disease, without killing the pathogen, and have been shown to cause asymptomatic carriage and transmission of lethal pathogens. Here we demonstrate that physiological defenses cooperate with the adaptive immune response to generate long-term asymptomatic carriage of the lethal enteric murine pathogen, *Citrobacter rodentium*. Asymptomatic carriage of genetically virulent *C. rodentium* provided immune resistance against subsequent infections. Host immune protection was dependent on systemic antibody responses and pathogen virulence behavior, rather than the recognition of specific virulent factor antigens. Finally, we demonstrate that an avirulent strain of *C. rodentium* in the field has background mutations in two genes that are important for LPS structure. Our work reveals novel insight into how asymptomatic infections can arise mechanistically with immune resistance, mediating exclusion of phenotypically virulent enteric pathogen to promote asymptomatic carriage.

## INTRODUCTION

For many infections, there is significant variation in host susceptibility to developing disease. For asymptomatic carriers, a pathogen can infect, replicate and transmit without causing clinical signs or symptoms of disease in the primary host (*1, 2*). Susceptible individuals that develop disease either recover after their sickness phase or ultimately continue to decline in health. A pathogen’s capability to cause sickness, aka “virulence”, depends on the damaging factors of the microbe as well as the host response to that pathogen. Much of the focus of understanding host susceptibility to infectious disease has focused on the host genetic makeup, metabolic or immune status, diet, and the microbiome that result in deteriorating health trajectories (*3–8*). We have relatively little understanding of how asymptomatic infections occur mechanistically or how they contribute to host defense and susceptibility to future infections. Because asymptomatic carriers contribute to infectious disease transmission and host-pathogen co-evolution, it is necessary to take a more holistic approach for our studies of infectious diseases and not solely focus on mechanisms resulting in sickness but also mechanisms of asymptomatic infections.

The host defense response, in part, dictates the heath trajectory a host will follow upon infection with a pathogen. Animals have evolved two distinct infection defense strategies that can be classified by their effects on pathogen fitness (*9*), i.e. antagonistic versus cooperative host defenses. Antagonistic defense mechanisms protect the host by having a negative impact on pathogen fitness. This includes mechanisms of immune resistance and nutritional immunity that operate to eliminate and kill pathogens (*10*). Antagonistic defenses also include behavioral avoidance mechanisms that reduce the risk of transmission (*11, 12*). Cooperative or physiological defense strategies mediate host adaptation to the infection, yielding an apparent truce between the host and pathogen (*13, 14*). This includes disease tolerance mechanisms that enable the host to withstand the presence of a pathogen by limiting physiological damage without killing the pathogen (*15–19*). Additionally, anti-virulence mechanisms are host physiological responses that can reduce the virulence behavior of the pathogen during colonization (*20, 21*). Since cooperative defenses sustain host health with a neutral to positive impact on pathogen fitness, it has been proposed that this defense strategy can promote asymptomatic carriage of pathogens (*9, 20*). Indeed, in a mouse model of infection with a lethal enteric pathogen, promoting anti-virulence defenses resulted in the persistent asymptomatic colonization of the pathogen (*20*). Studies to date have largely considered cooperative and antagonistic defenses as two independent defense strategies, though it is possible that these mechanisms may synergize to promote host survival and asymptomatic infections.

The mouse-specific attaching and effacing (A&E) pathogen, *Citrobacter rodentium*, causes infectious colitis, which mimics diseases caused by human-associated A&E pathogens, enterohemorrhagic *Escherichia coli* (EHEC) and enteropathogenic *E. coli* (EPEC) (*22–26*). Susceptibility to *C. rodentium* infection is dependent on the genetic background of the mice (*27, 28*). Whereas C57Bl/6 (B6) mice are highly resistant to *C. rodentium*, resulting in mild colonic inflammation and clearance of the pathogen by three weeks post-infection, *C. rodentium* causes severe and acute intestinal injury in C3H mice (*29, 30*). The ability of *C. rodentium* to cause disease requires virulence genes within the locus of enterocyte effacement (LEE) pathogenicity island (PAI) which encodes for a type 3 secretion system (T3SS) and effectors. The LEE PAI is regulated by the master virulence regulator, Ler, which is required for bacterial attachment and remodeling of the intestinal epithelium (*31, 32*). We previously investigated host metabolic adaptations to *C. rod*entium infection that regulate disease severity using the C3H model (*20*). In mice fed a normal diet, LEE virulence factor expression begins immediately after infection, with continued expression at the time of death around day nine post infection. We demonstrated that administration of an iron enriched diet led to activation of host anti-virulence defense mechanisms that involved transient insulin resistance and increased availability of glucose in the gut, which limited LEE virulence expression by the pathogen, resulting in asymptomatic, persistent carriage of the pathogen. This phenotypic attenuation persisted even after withdrawal of iron diet. This was followed by within-host evolution of genotypically attenuated *C. rodentium* characterized by non-functional mutations within the LEE PAI leading to an apparent commensalism between the host and *C. rodentium*.

In the current study, we used this dietary iron system to investigate whether immune resistance created in this setting is necessary for asymptomatic carriage of *C. rodentium*. We found that adaptive immunity was not necessary for the iron mediated anti-virulence mechanism, however, the adaptive immune response was necessary for the continued suppression of virulence and asymptomatic persistence of the pathogen.

Furthermore, asymptomatic carriage of genetically virulent but not genetically attenuated *C. rodentium* conferred protection from subsequent challenges with the parental virulent strain of *C. rodentium*. Virulence behavior of *C*. rodentium, but not virulence factor antigens, was necessary to induce protective immunity, leading to systemic, but not intestinal, antibody production. Finally, we demonstrate that an avirulent *ler* mutant of *C. rodentium* used in the field has secondary mutations in in LPS biosynthesis. Our work reveals novel insight into how asymptomatic infections can arise mechanistically and that physiological defenses cooperate with immune resistance to confer protection against lethal infections.

## RESULTS

### Adaptive immunity is required for asymptomatic carriage of *C. rodentium*

A schematic of our dietary iron model of *C. rodentium* asymptomatic infection (*20*) is shown in **Figure 1A**. We asked what host defense mechanisms are necessary for the continued suppression of *C. rodentium* virulence and asymptomatic carriage after removal of the iron diet. Specifically, we hypothesized that the adaptive immune system is necessary for asymptomatic carriage of *C. rodentium*. Thus, we generated immunodeficient SCID mice on the C3H/Snell background that are deficient for functional T and B cells (*SCID*^*-/-*^) and challenged them with wild-type *C. rodentium* in the presence of iron rich diet or control chow. *SCID*^*-/-*^ mice given dietary iron for two weeks had an initial survival advantage compared to both *SCID*^*+/+ or +/-*^ and *SCID*^*-/-*^ infected mice fed a control chow diet (**Figure 1B**). However 100% of the iron fed *SCID*^*-/-*^ mice eventually died ∼3 weeks post-infection (one week post-iron diet withdrawal), while iron fed *SCID*^*+/+ or +/-*^ with functional adaptive immunity survived and were protected from infection-induced weight loss (**Figure 1B-C, Supplemental Figure 1A**).

**Figure 1.**
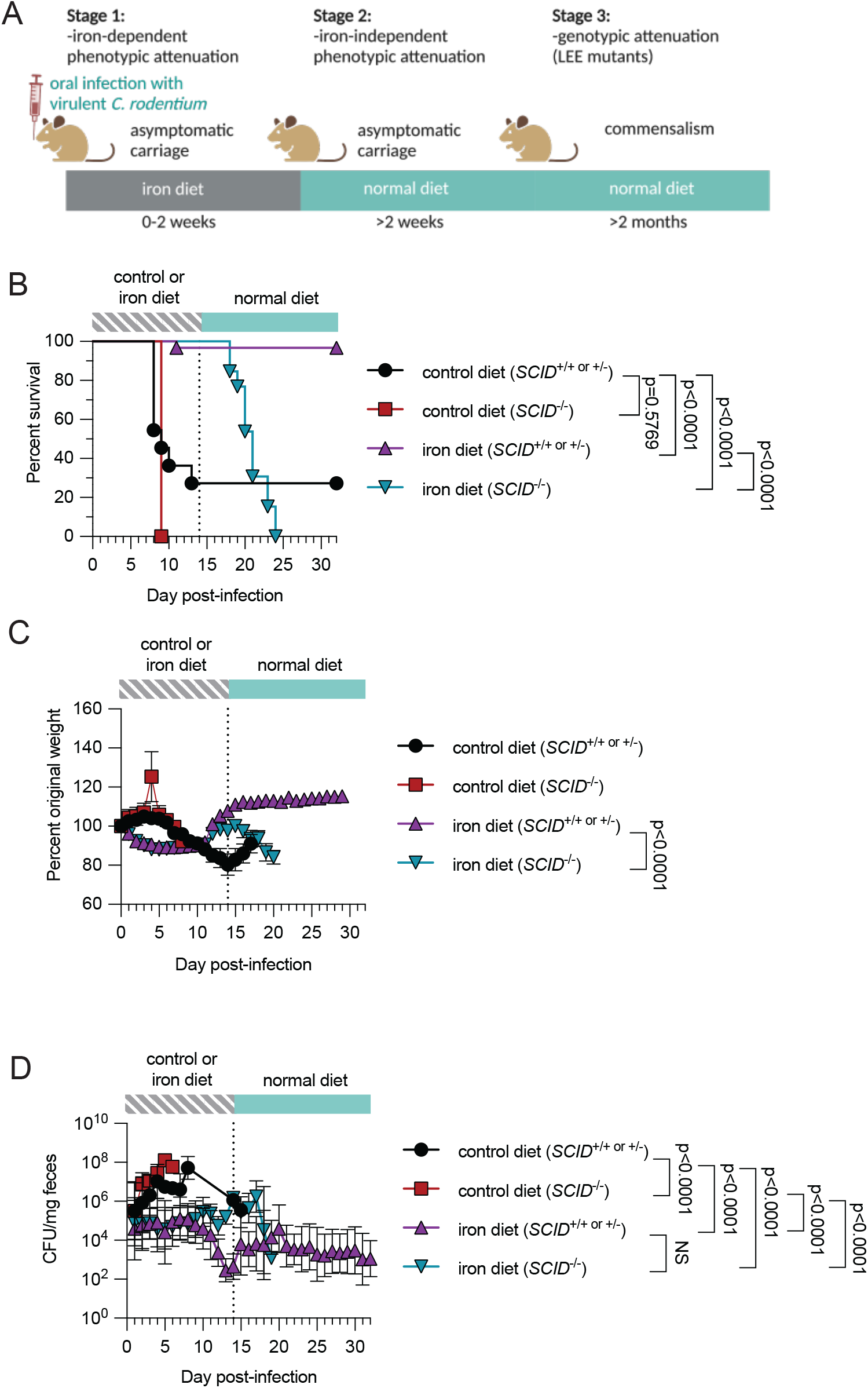
Adaptive immunity is required for asymptomatic carriage of *C. rodentium*. (A) Schematic showing the dietary iron mediated asymptomatic carriage of *C. rodentium* and subsequent evolution of attenuation (*20*). In Stage 1, dietary iron causes transient insulin resistance that is necessary to suppress expression of LEE PAI genes resulting in phenotypic attenuation. In Stage 2, the host continues to remain asymptomatically colonized with *C. rodentium* that is dependent on an iron-independent mechanism of phenotypic attenuation. In Stage 3, an apparent commensalism between the host and *C. rodentium* has formed due to within host evolution of *C. rodentium* that results in accumulation of mutations in LEE PAI genes. (B) survival, (C) percent original weight and (D) fecal shedding of immunodeficient C3H *SCID* mice (*SCID*^*-/*-^) or wild-type/heterozygous controls (*SCID*^*+/+ or +/*-^) infected with wild-type *C. rodentium*. At the time of infection, mice were fed *ad libitum* control or iron diet for two weeks before returning to normal vivarium chow. Control diet *SCID*^*+/+ or +/*-^ (n=11), control diet *SCID*^*-/*-^ (n=3), iron diet *SCID*^*+/+ or +/*-^ (n=31) and iron diet *SCID*^*-/*-^ (n=13). Data represent two biological replicates combined. (C) Error bars +/-SEM, (D) Geometric mean +/-geometric SD. Log rank analysis for survival and Two way ANOVA for pairwise comparisons.

Thus the adaptive immune response is not necessary for iron mediated protection from infection but is required for long-term survival during the iron independent phase. Examination of fecal *C. rodentium* levels over the course of the infection showed that there were comparable levels of pathogen in *SCID*^*+/+ or +/-*^ and *SCID*^*-/-*^ infected mice fed dietary iron at the onset of *SCID*^*-/-*^ death, and with no significant differences in pathogen burdens at any other time point assessed in in either the iron dependent or iron independent phase. This indicates that *SCID*^*-/-*^ mice have an impaired ability to adapt to the infected state (**Figure 1D and Supplemental Figure 1B**). Thus, the adaptive immune response is necessary for the continued suppression of virulence and asymptomatic carriage of an enteric pathogen in an iron-independent manner.

### Asymptomatic carriers are protected from subsequent challenges with *C. rodentium*

Next, we determined the consequences of asymptomatic carriage of a pathogen for host defense against subsequent challenges with the same pathogen. The experimental paradigm is depicted in **Figure 2A**. First, we challenged C3H/HeJ mice with an oral dose of PBS (mock 1°) or an oral dose of 7.5 × 10^8^ CFU of wild-type *C. rodentium* (wild-type 1°) which is normally a lethal infection in mice fed a normal diet (*20*) (**Figure 1B**). Mice were provided iron diet for two weeks to induce asymptomatic carriage of the pathogen, with fecal pathogen burdens peaking at 10^6^ CFU/mg feces (*20*) (**Figure 2B-C, Supplemental Figure 2A-C**). At 2 weeks post-infection with the primary challenge, the iron diet was removed and mice were returned to a normal chow diet. During this phase, pathogen burdens persisted at over 10^4^ CFU/mg feces (**Supplemental Figure 2A-B**). At 4 weeks post-infection with the primary challenge, we re-challenged the same mice with 7.5 × 10^8^ CFU of chloramphenicol (Cm) resistant wild-type *C. rodentium* (2° infection) to enable tracking of the secondary infection and kept mice on a normal chow diet. As expected, primary mock infected mice given dietary iron were highly susceptible to the wild-type secondary challenge, with all mice exhibiting severe weight loss and an inability to resist *C. rodentium* growth; eventually succumbing to the infection (**Figure 2B-D, Supplemental Figure 2C-D**). By contrast, 100% of mice that were originally challenged with wild-type *C. rodentium* for the primary round of infection (wild-type 1°) and given dietary iron were protected from weight loss and survived the secondary challenge with *C. rodentium* (**Figure 2B-C, Supplemental Figure 2C**). One possible model to explain this protection is that the host developed immune resistance against the secondary challenge to prevent colonization. Alternatively, following the primary infection, the host may sustain its anti-virulence response which suppresses virulence of the secondary *C. rodentium* infection. Such a scenario might occur through sustained metabolic/physiological changes or sustained changes in the microbiome (*33*). To distinguish between these two models, we quantified the levels of Cm-resistant *C. rodentium* that was being shed from primary mock- and wild-type-infected mice. In mock-infected mice, the levels of Cm-resistant *C. rodentium* reached 10^8^ CFU/mg feces by day 10 post-infection just prior to death. By contrast, there was a six log reduction in Cm-resistant *C. rodentium* being shed in the feces after the wild-type 1° with a low level persisting by three weeks post-infection (**Figure 2D, Supplemental Figure 2D**). Thus, asymptomatic carriage with *C. rodentium* seems to protect from subsequent lethal challenges via an antagonistic strategy, for example by heightening the host resistance defenses.

**Figure 2.**
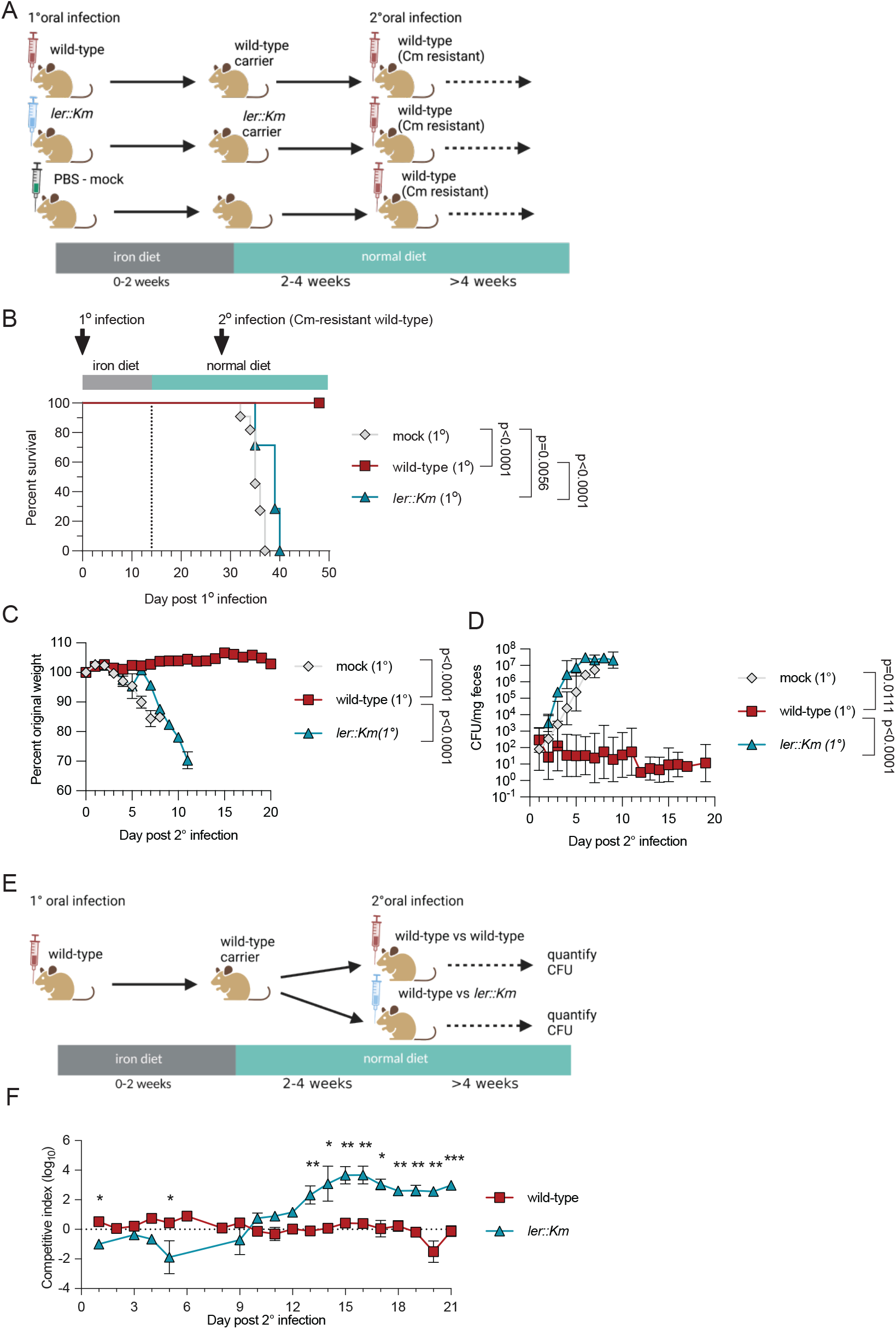
Asymptomatic carriers are protected from subsequent challenges with *C. rodentium*. (A) Schematic of secondary challenge experiments. C3H/HeJ mice were infected with of PBS (mock), or 7.5 × 10^8^ CFU of wild-type or *ler::Km* and fed iron diet for two weeks. At 4 weeks post-infection mice were re-challenged with 7.5 × 10^8^ CFU of Cm-resistance wild-type *C. rodentium*. (B) Survival curve, (C) percent original weight post secondary infection, (D) fecal shedding of Cm-resistant *C. rodentium*. n = 7 to 19 mice per condition. Data represent three biological replicates combined. (E) Schematic of competition experiments. C3H/HeJ mice were infected with 7.5 × 10^8^ CFU wild-type *C. rodentium*. Mice were fed iron diet between zero and two weeks post-infection. At 4 weeks post-infection mice were challenged with different antibiotically tagged wild-type and *ler::Km* strains at a 1:1 ratio. (F) Competitive index scores over time calculated as wild-type or *ler::Km* over wild-type. n = 5 mice per condition. Data represent one biological replicate. Error bars are +/-SEM for (C) and (F). Geometric mean +/-geometric SD for (D). \**p* < 0.05, \*\**p* < 0.01, \*\*\**p* < 0.001. Log rank analysis for survival. Unpaired t-test or Two way ANOVA for pairwise comparisons.

We next investigated how the host can accommodate persistent colonization with *C. rodentium* that are phenotypically attenuated, but can mount an effective resistance response against the secondary challenge. We hypothesized that the resistance response can selectively exclude phenotypically virulent pathogen. We tested this idea with a competition experiment as shown in **Figure 2E**. We engineered different antibiotic tagged versions of wild-type *C. rodentium* and obtained an attenuated *C. rodentium* strain, *ler::Km*, which encodes a mutation in the master virulence regulator in *C. rodentium* (*31*). *C. rodentium* strains lacking *ler* do not cause disease and are completely attenuated in mice (*31*). Compared to mice that are singly infected with the wildtype strain, mice singly infected with the *ler::Km* strain show reduced colonization when fed iron as indicated by fecal shedding levels. However after two weeks post-infection when mice are returned to normal chow, the colonization levels are comparable to that found in mice singly infected with the wild type strain (**Supplemental Figure 2A-B**). *In vitro*, growth of the wild-type and attenuated strain were comparable, demonstrating no inherent growth advantages exist (**Supplemental Figure 2E**). We infected C3H/HeJ mice with a 7.5 × 10^8^ CFU dose of wild-type *C. rodentium* and provided iron chow. After 2 weeks, iron chow was removed, and mice were returned to their normal chow diet. At four weeks post primary challenge, we rechallenged mice with a 1:1 dose of wild-type and the attenuated *ler::Km C. rodentium* and continued to feed on normal chow. Although the attenuated *C. rodentium* strain displayed an initial competitive disadvantage, it quickly outcompeted the wild-type strain of *C. rodentium* after one week post-infection (**Figure 2F and Supplemental Figure 2F**). Importantly, the competitive advantage of the *ler::Km* strain was not observed in previously unchallenged mice (**Supplemental Figure 2G-H**). These data demonstrate that mice asymptomatically colonized with wild-type *C. rodentium* can selectively exclude subsequent challenges with virulent but not LEE-attenuated *C. rodentium*.

Although dietary iron limits the expression of the LEE virulence program to prevent detectable clinical and pathological effects of a primary *C. rodentium* infection on the host (*20*), because there is selective exclusion of virulent *C. rodentium* in response to the secondary challenge, we considered the hypothesis that virulence-competent *C. rodentium* is required in the primary infection to mediate resistance defenses against subsequent challenges with the same pathogen. To test this, we followed the experimental paradigm in **Figure 2A**, and challenged mice on iron diet with wild type *C. rodentium* or *ler::Km* strain for their primary challenge. In C3H/HeJ mice fed dietary iron, the *ler::Km* strain is not completely cleared, as indicated by persistent fecal shedding post primary infection (**Supplemental Figure 2A**). After two weeks, iron diet was removed and the mice were returned to control chow. At 4 weeks post-infection mice were re-challenged with a lethal dose of wild-type Cm-resistant *C. rodentium*. 100% of mice that received the primary challenge with virulent *C. rodentium* were protected from the secondary challenge but mice that received *ler::Km* for the primary challenge were susceptible to the wild-type secondary challenge (**Figure 2B-C, Supplemental Figure 2C**).

Similar to mock 1° mice, *ler::Km* 1° mice susceptibility was associated with an inability to control Cm-resistant *C. rodentium* levels (**Figure 2D, Supplemental Figure 2D**). We also repeated these protection experiments with a *C. rodentium* isolate naturally missing the LEE PAI (herein called ∆LEE, isolate #9 from *20*). Like *ler::Km*, primary infection with ∆LEE strain also did not confer protection from secondary infection by wild-type *C. rodentium* (**Supplemental Figure 2I**). Together these data show that the presence of LEE encoded virulence factors during the primary challenge in the asymptomatic host are required to mediate protective immunity against subsequent challenges of virulent *C. rodentium*.

### *C. rodentium* specific antibodies do not distinguish between wild-type and Ler/LEE deficient strains

Our experimental data thus far demonstrate that 1) adaptive immunity is necessary for asymptomatic colonization and long-term survival of the host in the iron-independent phase; 2) mice develop a form of protective memory against subsequent challenges through selective exclusion of phenotypically virulent *C. rodentium;* and 3) development of this memory is dependent on LEE-encoded virulence in the iron dependent phase. We reasoned that the mechanism of adaptive immune mediated suppression of virulence may require antibody mediated defenses and that this could be extended to provide protection against subsequent challenges with virulent *C. rodentium*. After *C. rodentium* oral infection, mice fed dietary iron for two weeks developed robust serum IgG and IgM, as well as intestinal IgA responses against wild-type *C. rodentium* (**Figure 3A-B, Supplemental Figure 3A-B**). Unlike previous reports (*34*), we were unable to detect *C. rodentium* specific IgG in the gut of C3H or B6 mice (**Figure 3A, Supplemental Figure 3A&C**). As expected, *SCID*^*-/-*^ or mock infected mice did not develop *C. rodentium* specific antibodies (**Figure 3A-B, Supplemental Figure 3A-D**). Finally, and consistent with a role for antibody mediated defense against *C. rodentium* (*35–40*), we found that muMT (B6 background) mice, which are deficient for mature B cells, were highly susceptible to an oral challenge with wild-type *C. rodentium* infection despite feeding of dietary iron (**Figure 3C-D, Supplemental Figure 3E**).

**Figure 3.**
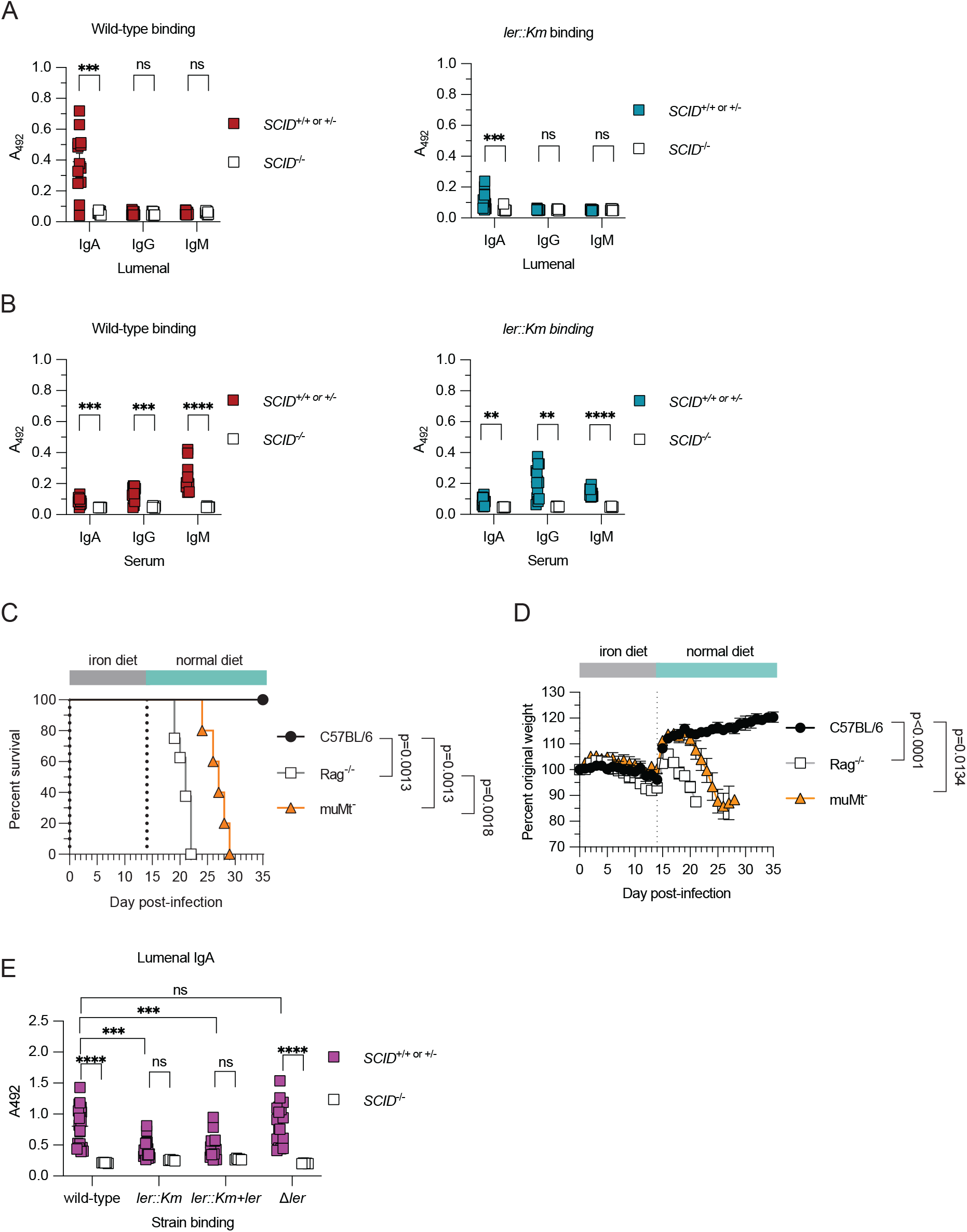
*C. rodentium* specific antibodies do not distinguish between wild-type and Ler/LEE deficient strains. Whole bacteria ELISAs quantifying (A) lumenal or (B) serum IgA, IgG or IgM antibodies binding to wild-type or *ler::Km C. rodentium*. n = 6 to 13 mice per condition. Data represent one biological replicate. (C) Survival and (D) percent original weight of wild-type, *Rag*^*-/*-^, and *muMt*^*-*^ mice infected with 7.5 × 10^8^ CFU wild-type *C. rodentium*. Mice were fed iron diet for two weeks and then placed on normal diet for the remainder of the experiment. n = 5 to 8 mice per condition. Data represent one biological replicate. (E) Whole bacterial ELISAs quantifying lumenal IgA binding against wild-type, *ler::Km, ler::Km+ler*, or *∆ler*. n = 5 to 19 mice samples per condition. Data represent one biological replicate. Lumenal and serum samples were collected from wild-type *C. rodentium* infected C3H SCID mice fed iron diet for two weeks post-infection. Mean +/-SEM. Unpaired t-test, Mann Whitney test or One-Way ANOVA with post Tukey test or Two way ANOVA performed for pairwise comparisons. Log rank analysis for survival. \**p* < 0.05, \*\**p* < 0.01, \*\*\**p* < 0.001, \*\*\*\**p* < 0.0001.

It has been previously reported that lumenal IgG *C. rodentium* specific antibodies bind to LEE encoded virulence factors and that this is necessary for the selective exclusion of virulent *C. rodentium* in the gut (*34*). We detected *C. rodentium* specific lumenal IgA, rather than lumenal IgG in our dietary iron system. Because we found that resistance against subsequent challenges of *C. rodentium* requires LEE genes in the primary infection and because previously challenged mice can selectively eliminate virulent *C. rodentium* in our dietary iron model, we posited that *C. rodentium* specific lumenal IgA antibodies were recognizing LEE encoded virulence factors in our system. From our antibody binding assays, we consistently saw that lumenal IgA isolated from wild-type *C. rodentium* infected mice fed dietary iron had diminished recognition against *ler::Km*, though binding were not completely ablated (**Figure 3A** and **Supplemental Figure 3A&C**). We also found that serum IgM antibodies isolated from wild-type *C. rodentium* infected mice fed dietary iron had diminished recognition against *ler::Km*, though binding were not completely ablated (**Figure 3B** and **Supplemental Figure 3B&D**). Additionally, we found that serum IgG bound to *ler::Km* strain as well as to wild type *C. rodentium* (**Figure 3B** and **Supplemental Figure 3B & D**) rather than a diminished response as previously reported for *Citrobacter* lacking *ler* (*34*). To determine if the reduction in *C. rodentium* lumenal IgA was due to the loss of *ler* and LEE virulence factor expression, we complemented the *ler* mutation in the *ler::Km* strain. Although complementation partially rescued growth of the strain in mice (**Supplemental Figure 3F-G**), complementation did not restore gut IgA antibody binding to the *ler::Km* strain (**Figure 3E**). We also examined lumenal IgA antibody binding to the ∆LEE strain. Consistent with our complementation studies, ∆LEE was still recognized by lumenal IgA at levels comparable to the wild-type strain (**Supplemental Figure 3H)**. Furthermore, we constructed a clean in-frame detection of *ler* named ∆*ler* and tested it for antibody binding. The Δ*ler* strain did not evade lumenal IgA or serum IgG antibody binding (**Figure 3E** and **Supplemental Figure 3I**) like the *ler::Km* strain. Together, these data suggest that discrimination between wild-type and attenuated *C. rodentium* by antibodies is not dependent on *ler* or *ler* regulated LEE encoded virulence factors.

### LPS is necessary for recognition by *C. rodentium* specific antibodies

Because we found intestinal IgA antibody binding was reduced with the *ler::Km* strain yet not restored with the *ler::Km+ler* complemented strain, it suggests that there are secondary mutations in the *ler::Km* strain resulting in antibody avoidance. We used this strain as a tool to investigate the IgA dependent antibody recognition of *C. rodentium*. We sequenced the genomes of wild-type, *ler::Km*, and Δ*ler* strains, performed *de novo* hybrid genome assemblies, and identified secondary mutations in the *ler::Km* strain. Mutational analysis of the *ler::Km* genome identified multiple IS102 transposon duplications and insertions in ROD_21691 and ROD_41941, likely inactivating them (**Supplemental Figure 4A, Supplemental Table 1**). Consistent with the genome analysis, PCR amplifications of those genes in *ler::Km* showed PCR products +1000bp greater than in either wild-type (**Supplemental Figure 4B**). ROD_21691 and ROD_41941 are putative glycosytransferase and lipopolysaccharide (LPS) acetylglucosaminyltransferases, respectively, and herein renamed *wfaP* and *rfaK* for their homology to *E. coli* LPS synthesis genes **(Supplemental Figure 4C-E)**. WfaP is a predicted glucosyltransferase responsible for polymerization of O-antigen of LPS (*41*), whereas RfaK is essential to complete synthesis of the core subunit of LPS (*42*).

To examine the role of the *rfaK* and *wfaP* transposon mutations on LPS structure and antibody recognition in the *ler::Km* strain, we constructed single and double *rfaK* and *wfaP* complements. LPS extractions and LPS blots revealed that *ler::Km* but not wild-type or our newly constructed Δ*ler* mutant is defective in LPS assembly (**Figure 4A**). Importantly, double complementation of *rfaK* and *wfaP* were required to restore LPS in the *ler::Km* strain (**Figure 4A**). This demonstrates that through some unknown events, the *ler::Km* strain had acquired two inactivating mutations within the same LPS biosynthesis pathway. We next asked whether these mutations were responsible for loss of antibody binding of the *ler:Km* strain. Indeed complementation of both the *rfaK* and *wfaP* mutations in *ler::Km* restored *C. rodentium* specific IgA recognition (**Figure 4B**). Interestingly *ler::Km* had partially enhanced, although not significant, IgG binding compared to wild-type, but binding was diminished when complemented with *rfaK* and *wfaP* (**Figure 4C**). This suggests that *C. rodentium* LPS may partially mask *C. rodentium* from IgG recognition. Finally, we generated clean in-frame deletions of *rfaK* and *wfaP* on the wild-type *C. rodentium* background. As predicted both ∆*rfaK* and ∆*wfaP* single gene knock out strains were defective in LPS synthesis (**Figure 4D**) and complementation restored full LPS synthesis (**Figure 4D**). When we examined antibody binding by whole bacterial ELISAs, we found that individual LPS mutations disrupted IgA binding to *C. rodentium* (**Figure 4E-F**). Together these data demonstrate that *C. rodentium* infection with our dietary iron model results in the generation of pathogen specific antibodies that require LPS and not *ler*-dependent antigens for binding.

**Figure 4.**
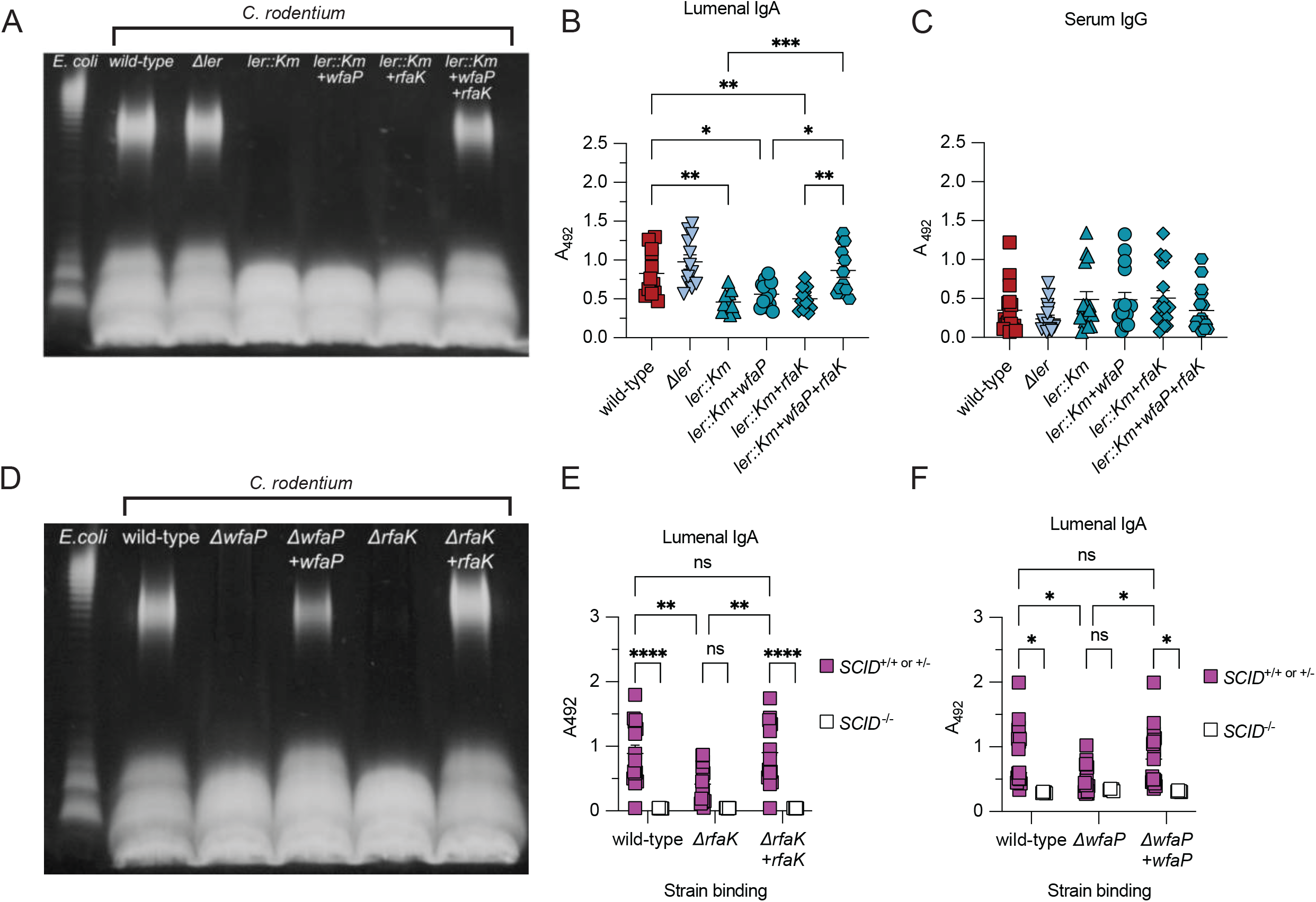
LPS, not ler, is recognized by *C. rodentium* specific antibodies. (A) LPS quantification of *E. coli*, wild-type, ∆*ler, ler::Km, ler::Km+wfaP, ler::Km+rfaK*, and *ler::Km+wfaP+rfaK*. (B-C) Whole bacteria ELISA quantifying (B) lumenal IgA or (C) serum IgG binding against wild-type, ∆*ler, ler::Km, ler::Km+wfaP, ler::Km+rfaK*, and *ler::Km+wfaP+rfaK*. n = 12 samples per condition for (B) and n = 16 samples per condition for (C). Data represent one biological replicate. (D) LPS quantification of *E. coli*, wild-type, ∆*wfaP, ∆wfaP+wfaP, ∆rfaK, ∆rfaK+rfaK*. (E-F) Whole bacteria ELISA quantifying (E) lumenal IgA or (F) serum IgG binding against wild-type, ∆*wfaP, ∆wfaP+wfaP* (E), *∆rfaK*, and *∆rfaK+rfaK* (F). n = 5 to 19 samples per condition. Data represent one biological replicate. Lumenal and serum samples were collected from wild-type *C. rodentium* infected C3H SCID mice fed iron diet for two weeks post-infection. Statistical significance was calculated using One Way ANOVA with post Tukey test, Kruskal Wallis with post Dunn’s test or two-way ANOVA. \**p* < 0.05, \*\**p* < 0.01, \*\*\**p* < 0.001, \*\*\*\**p* < 0.0001

### Adaptive immunity regulates Ler and not LPS dependent virulence of *C. rodentium*

We next tested the hypothesis that the adaptive immune system is necessary to control LPS-dependent virulence, rather Ler/LEE dependent virulence. We first tested the importance of LPS for *C. rodentium* virulence *in vitro* by determining if the LPS strains were defective in pedestal formation, a requirement for *in vivo* virulence (*43*), on HeLa cells. Both ∆*wfaP* and ∆*rfaK* induced pedestals but not attenuated controls, ∆*ler* or ∆*tir*, a mutant lacking the translocated intimin receptor (*tir*) required for cell attachment (*43*) (**Figure 5A**). *In vivo*, infection of mice fed a normal chow diet with the ∆*wfaP* strain resulted in a mild delay in host weight loss and death kinetics compared to mice infected with the wild-type strain of *C. rodentium*. Analysis of pathogen burdens revealed no significant differences in peak pathogen burdens, although there was a delay in rate at which peak levels were reached in the ∆*wfaP* infected mice (**Figure 5B-D, Supplemental Figure 5A&B**).

**Figure 5.**
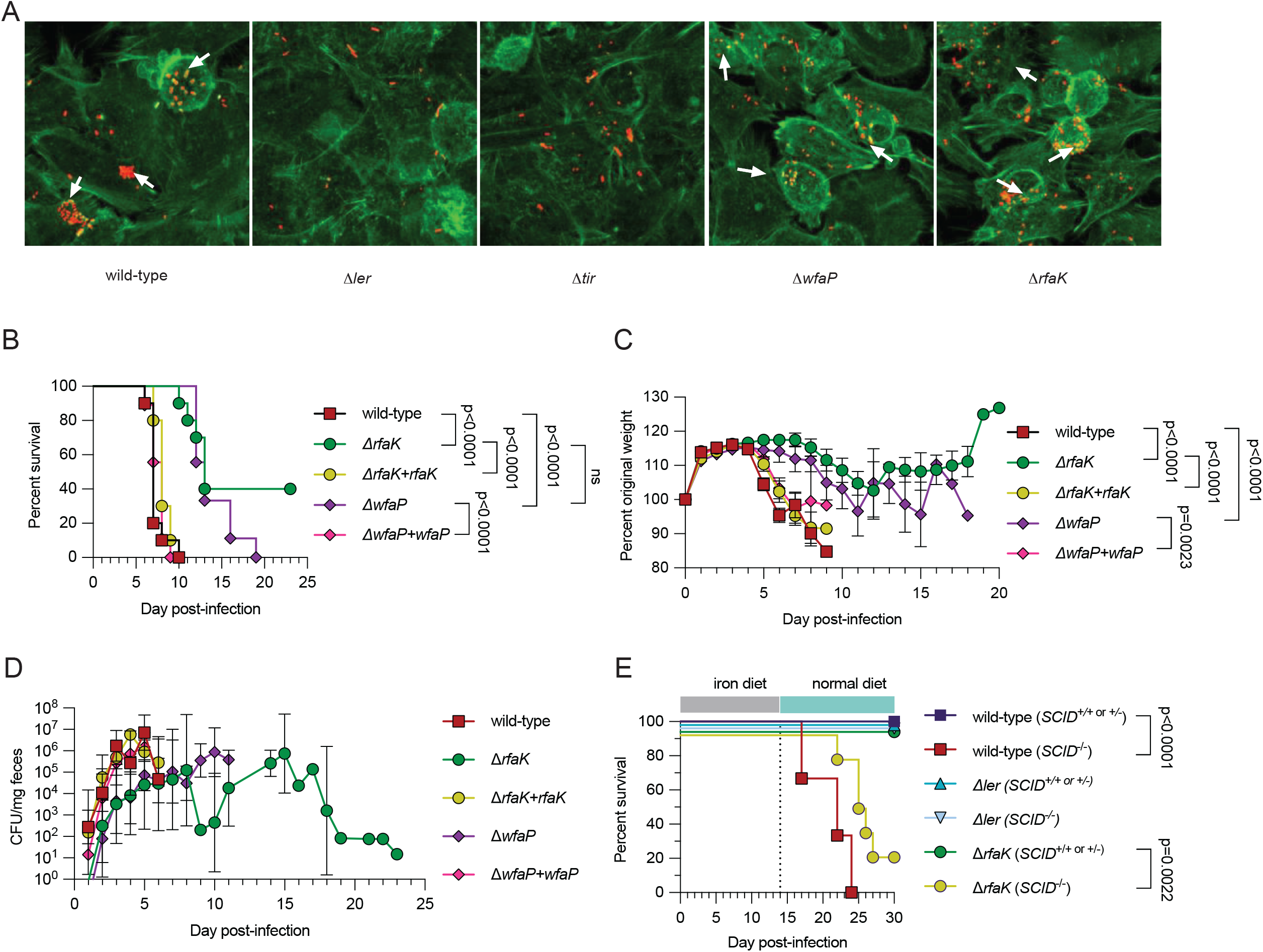
LPS promotes virulence of *C. rodentium*. (A) Representative images of FAS assay of wild-type, ∆*ler, ∆tir, ∆wfaP*, and *∆rfaK* mutants attaching to HeLa cells. Green signifies host cell actin and red signifies *C. rodentium*. White arrows denote foci of actin associated with bacteria. (B-D) C3H mice were infected with 7.5×10^8^ CFU of wild-type, ∆*rfaK, ∆wfaP*, and complement strains. (B) Survival, (C) percent original weight, and (D) fecal shedding. n = 9-10 mice per condition. Data represent one biological replicate. (E) *SCID*^*+/+or+/-*^ and *SCID*^*-/-*^ mice were infected with wild-type, ∆*ler*, or *∆rfaK* and given iron diet for two weeks after which mice were switched back to their normal chow diet. Survival was determined. Wild-type (*SCID*^*+/+ or +/*-^) = 5 mice, wild-type (*SCID*^*-/-*^) = 3 mice, *Δler* (*SCID*^*+/+ or +/-*^) = 2 mice, *Δler* (*SCID*^*-/-*^) = 3 mice, Δ*rfaK* (*SCID*^*+/+ or +/-*^) = 9 mice, Δ*rfaK* (*SCID*^*-/-*^) = 7 mice. Data represent two biological replicates combined. Log rank analysis for survival. Two way ANOVA for pairwise comparisons. Error bars indicate +/-SEM for weight curves and Geometric mean +/-geometric SD for CFU analyses.

Complementation of the ∆*wfaP* strain with a wild-type copy of the gene restored virulence of the strain (**Figure 5B-D, Supplemental Figure 5A**). Infection with the ∆*rfaK* strain exhibited a slightly greater attenuated phenotype, with only 60% of infected mice succumbing to the infection and also protection from clinical signs of disease (**Figure 5B-D, Supplemental Figure 5A&B**). Analysis of the fecal pathogen burdens revealed that the ∆*rfaK* strain reached comparable peak levels as the virulent strains, however the rate at which peak levels were reached was slower, indicating that LPS is necessary for *C. rodentium* virulence *in vivo*. Thus, with respect to the wild type strain, *C. rodentium* defective in LPS is mildly attenuated, yet is still capable of causing a lethal infection in mice under normal chow conditions.

We next tested the effects of LPS deficiency for *C. rodentium* virulence in SCID deficient mice with our dietary iron model. Similar to mice infected with the wildtype *C. rodentium* strain, a two week course of dietary iron was sufficient to protect *SCID*^*+/- or +/-*^ mice from lethality and weight loss when infected with the Δ*rfaK* strain (**Figure 5E, Supplemental Figure 5C-D**). By contrast *SCID*^*-/-*^ mice succumb to infection with Δ*rfaK* despite receiving the two week course of dietary iron (**Figure 5E, Supplemental Figure 5C-D**). Instead, we found that infection of *SCID*^*-/-*^ mice with the Δ*ler* strain is attenuated (**Figure 5E, Supplemental Figure 5C-D**). Taken together, our data demonstrate that the adaptive immune and humoral response is necessary for defense against Ler-dependent virulence, however this does not require antibody binding to Ler encoded antigen as has been previously suggested.

### Pathogen behavior is necessary for protection of asymptomatic carriers with subsequent infections

Mice fed dietary iron and infected with the *ler::Km* strain of *C. rodentium* are not protected from subsequent challenges with the parental virulent strain of the pathogen (**Figure 2**). While our studies with the ΔLEE strain (**Supplemental 2I**) suggests that LEE virulence factors are required during the primary infection for the protection of asymptomatic carriers against subsequent challenges, because we found the *ler::Km* to have background mutations in genes important for LPS synthesis, we wanted to rigorously test a role for LPS in addition to Ler for mediating this protective effect using our clean deletion strains. First we tested the importance of LPS and Ler for the generation of *C. rodentium* specific antibodies during infection. *SCID*^*+/+ or +/-*^ and *SCID*^*-/-*^ mice were orally infected with wild-type, the LPS defective Δ*rfaK*, or the Δ*ler* strain and given iron chow. We then collected lumenal and serum antibodies from mice at 14 days post-infection and tested their ability to bind specifically to wild type *C. rodentium*. Mice infected with Δ*rfaK* were unable to generate gut IgA antibodies that recognize wild-type *C. rodentium* (**Figure 6A**); however, ∆*rfaK* infected mice did generate *C. rodentium* specific serum IgG antibodies (**Figure 6B**). Mice infected with Δ*ler* did not generate lumenal IgA or serum IgG antibodies that bind to wild type *C. rodentium* (**Figure 6A-B**).

**Figure 6.**
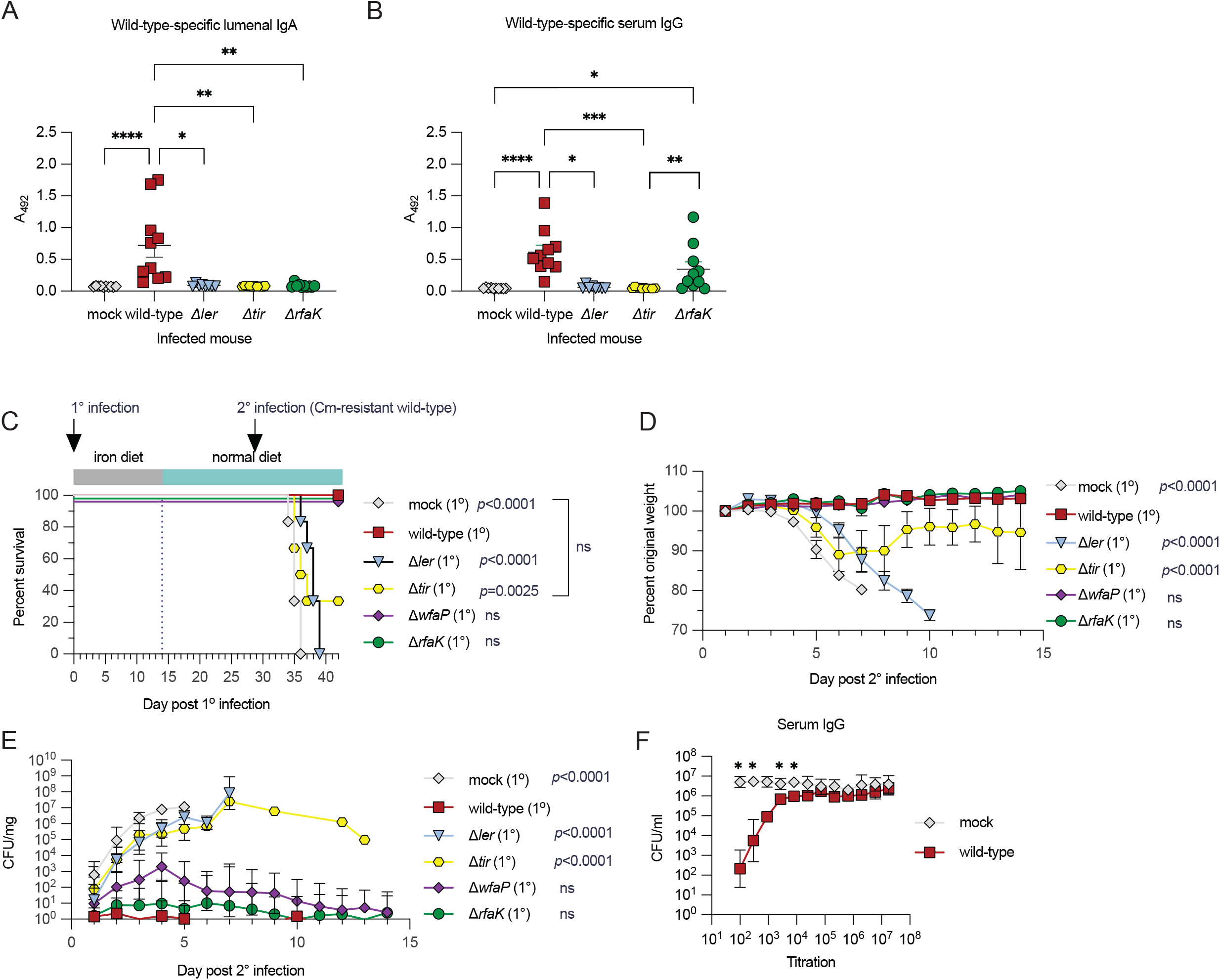
Pathogen behavior is necessary for protection of asymptomatic carriers with subsequent infections. (A-B) Whole bacteria ELISAs quantifying (A) lumenal IgA or (B) serum IgG against wild-type *C. rodentium*. Antibody samples were collected at two weeks post-infection from C3H mice infected with PBS (mock), wild-type, ∆*ler, ∆tir*, or *∆rfaK* and fed iron diet. n = 8-10 mice per condition. Data represent one biological replicate. (C-E) Secondary challenge experiments were performed as depicted in Figure 2A but with primary infections of PBS (mock), wild-type, ∆*ler, ∆tir, ∆wfaP, or ∆rfaK* and secondary infections with Cm-resistant wild-type *C. rodentium*. (C) Survival, (D) percent original weight, and (E) fecal shedding were monitored. n = 6-10 mice per condition. Data represent two biological replicates combined. (F) Complement bactericidal assay using serum from mock infected or wild-type infected *C. rodentium* mice fed iron. Serum was collected at 14 days post-infection and killing was tested on wild-type *C. rodentium*. n = 3 mice per condition. One way ANOVA with post Tukey test, Kruskal Wallis with post Dunns test, Two way ANOVA or unpaired t test for pairwise comparisons. Log rank analysis for survival. Error bars are +/-SEM for (D) and Geometric mean +/-geometric SD for (E-F). \**p* < 0.05, \*\**p* < 0.01, \*\*\**p* < 0.001, \*\*\*\**p* < 0.0001. P values in (C-E) are shown condition vs wild type.

To determine the consequences of LPS and ler deficiency for an asymptomatic carriers’ ability to protect against subsequent *C. rodentium* challenges, we infected C3H mice with wild-type, *∆ler*, ∆*wfaP* or ∆*rfaK* strains or treated with PBS for mock infection, and gave iron diet for two weeks. At four weeks post-infection with the primary challenges, we re-challenged the mice with a lethal dose of wild-type *C. rodentium*. Similar to *ler::Km* (**Figure 2B**), ∆*ler* also did not provide protective immunity (**Figure 6C-E**). Interestingly, mice immunized with the LPS strains were completely protected from secondary challenge with the wild-type strain demonstrating that the presence of LPS is not necessary for development of resistance defenses against subsequent virulent challenges (**Figure 6C-E, Supplemental Figure 6A-B**). These data also demonstrate that serum IgG responses correlate with potential for protective immunity, suggesting that serum IgG is important for defense against *C. rodentium* while lumenal IgA is dispensable. Indeed, in a complement bactericidal assay, serum from iron fed mice and infected with wild type *C. rodentium* kills the pathogen (**Figure 6F, Supplemental Figure 6C**). By contrast, unlike *SCID*^*-/-*^ and muMt mice, *IgA*^*-/-*^ mice are completely protected from infection with wild type *C. rodentium* (**Supplemental Figure 6D**). Thus, ler and LEE encoded virulence factors are necessary for resistance against secondary challenges in asymptomatic carriers of *C. rodentium*, via the induction of serum IgG responses.

There are several models to explain how *ler* and LEE encoded virulence factors can mediate the generation of serum IgG and protection against secondary *C. rodentium* challenges. The first model is that the host recognizes specific virulence factor antigens that are expressed on virulent *C. rodentium* but not the attenuated strains that are required for the generation of serum IgG. A second model is that LEE encoded virulence factors enable the pathogen to invade the host niche, which is necessary for pathogen recognition and the generation of serum IgG. Our data argues against LEE specific antigens as the sole mediator of protection and so we consider the later model. Both the Δ*ler* strain and the Δ*tir* strain are defective in attachment/invasion (*31, 43*) (**Figure 5A**). However, while the Δ*ler* strain is defective in the expression of virulence factors within the LEE pathogenicity island (**Supplemental Figure 6E-F**), the Δ*tir* strain is not defective in expression of LEE virulence genes (*31*) (**Supplemental Figure 6E-F**). We used these strains to discriminate against the requirement for LEE virulence factors and pathogen behavior for regulation of *C. rodentium* virulence and protection against secondary challenges. Mice that received Δ*tir* for their primary challenge were completely susceptible to secondary infections (**Figure 6C-E, Supplemental Figure 6A-B**). Furthermore, mice infected with the Δ*tir* strain fail to produce *C. rodentium* specific serum IgG or lumenal IgA antibodies (**Figure 6A-B**). Taken together our data demonstrate that virulence activity or behavior is a prerequisite for long-term protection and asymptomatic carriage of phenotypically attenuated *C. rodentium* (**Figure 7**).

**Figure 7.**
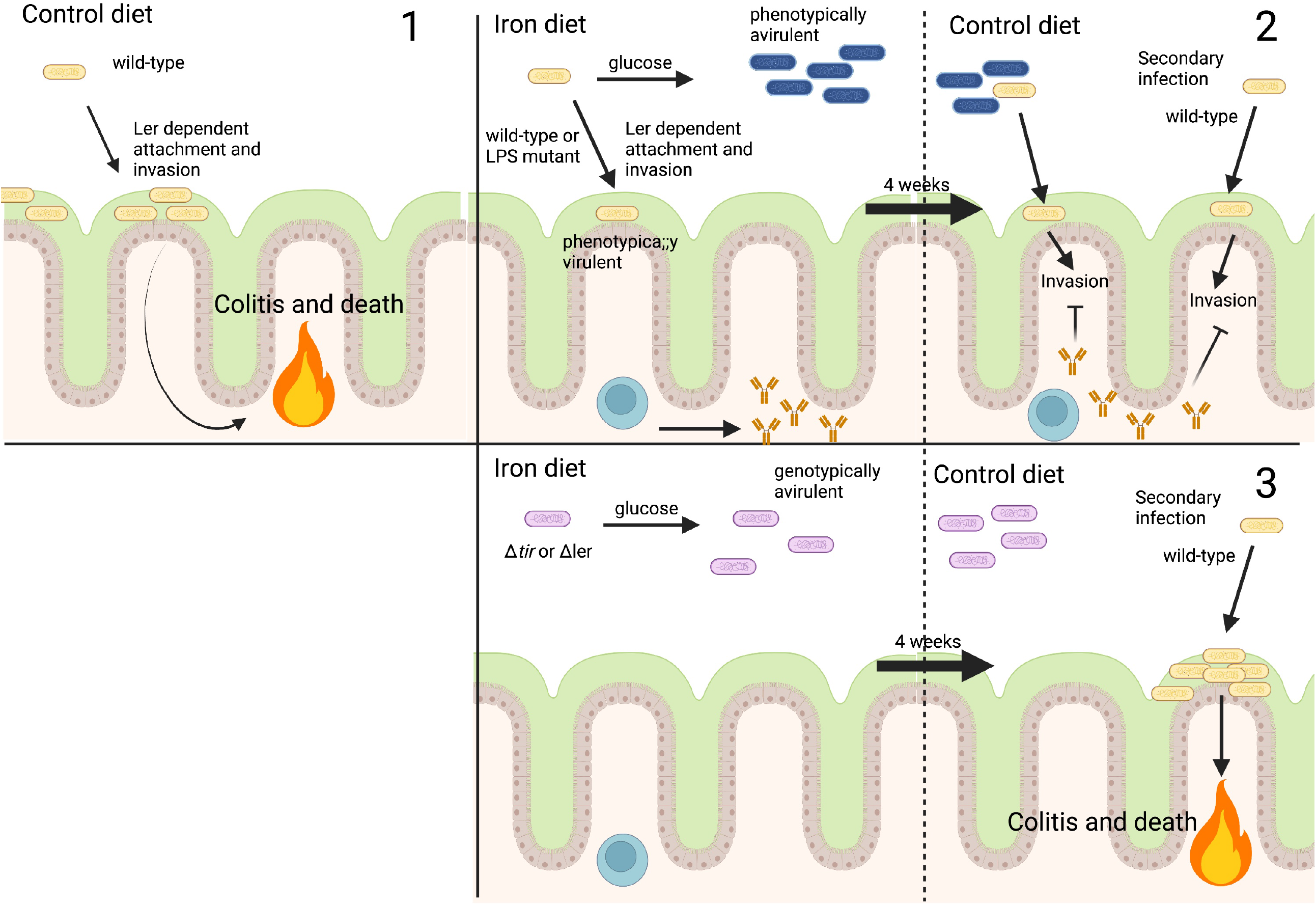
Adaptive immunity and pathogen behavior drive asymptomatic carriage. Model of how iron induced metabolic adaptations promote phenotypic attenuation of *C. rodentium*, development of adaptive immunity, long-term asymptomatic carriage, and protection from subsequent infection. LEE pathogenicity is required to prime the immune system and promote immune protection. 1. Under control diet conditions, wildtype *C. rodentium* expresses Ler and LEE virulence factors facilitating attachment, invasion, colitis and death. 2. Under dietary iron conditions, there is an increase in glucose availability in the intestine (*20*), which suppresses expression of LEE encoded virulence factors resulting in a largely phenotypically avirulent population of *C. rodentium* that remains lumenally bound. A small proportion of the pathogen population expresses LEE virulence factors, attaches to the epithelium and induces systemic *C. rodentium* IgG, which is necessary for the selective exclusion of phenotypically virulent *C. rodentium* that arises after iron withdrawal as well as protection from subsequent challenges with wild-type pathogen. 3. When infected with the Δ*ler* or Δ*tir* strain, there is no phenotypically virulent pathogen to adhere to the intestinal epithelium and induce a systemic IgG response. When animals are challenged with a secondary wild-type *C. rodentium* infection they are susceptible due to the lack of *C. rodentium* specific IgG.

## DISCUSSION

Asymptomatic infections contribute to infectious disease transmission and host-pathogen co-evolution. However, while we have made great advances in our mechanistic understanding of how pathogens cause disease, we have relatively no understanding of the mechanisms that facilitate the ability of pathogens to infect and replicate inside the host niche without causing disease. We previously demonstrated that dietary interventions can yield asymptomatic infections with the enteric pathogen *C. rodentium*, by promoting an anti-virulence defense mechanism that prevents engagement of the pathogens virulence program without affecting its ability to colonize or replicate in the host (*20*). This phenotypic attenuation lasted for months after the removal of the dietary intervention, prompting us to ask whether host adaptive immunity contributes to the regulation of virulence and asymptomatic persistence in our model system. Consistent with the current knowledge about *C. rodentium* immunity (*35, 38, 39, 44*), we found long-term protection against *C. rodentium* to require a functional adaptive immune system. Our data supports a model whereby the host transitions from anti-virulence to immune resistance defenses to sustain asymptomatic persistence by selectively excluding phenotypically virulent pathogen, thereby, allowing phenotypically avirulent pathogen to persist (*45*). We suggest this is dependent on pathogen behavior and systemic IgG responses. In some contexts, it is logical to conceptualize cooperative defenses as a strategy to buy the host time for their adaptive immune response to kick in and clear the infection. Collectively our studies demonstrate that indeed there is a cooperation between physiological defenses and adaptive immune resistance in defense against a lethal bacterial infection, and that this can unexpectedly yield sustainment of an asymptomatic infection.

Our data from the current study support a model that is mechanistically distinct from previous studies examining the interplay between the adaptive immune response, virulent and avirulent *C. rodentium*. Using a C57Bl/6 model, Kamada et al. (*34*) also reported that there were subpopulations in *C. rodentium* infected mice that were phenotypically virulent and avirulent, and that the adaptive immune response was necessary for the selective exclusion of virulent *C. rodentium* in the intestine. The authors proposed that gut IgG, and not IgA, binding of LEE encoded virulence factor antigens that were expressed on virulent but not avirulent *C. rodentium* facilitated neutrophil mediated elimination. In our studies we were unable to detect *C. rodentium* specific IgG in the gut and could only detect *C. rodentium* specific IgA in the intestines of infected mice. We report that *C. rodentium* specific gut IgA binding is not dependent on the presence of Ler/LEE virulence factors, and instead binding is largely dependent on LPS. In any case, we demonstrate that gut IgA binding of LPS is not necessary for the selective exclusion of virulent *C. rodentium*. Instead, our data show that systemic *C. rodentium* specific IgG mediates the selective exclusion of virulent *C. rodentium*. This is in agreement with previous studies from multiple groups demonstrating the importance of systemic IgG in host resistance against *C. rodentium* (*34, 35*). Using the Δ*tir* mutant, we further demonstrate that virulent behavior (attachment/pedestal formation) mediated by LEE encoded virulence factors rather than recognition of virulence factor antigens is what is necessary for selective exclusion of phenotypically virulent *C. rodentium* in the host niche. Some possible explanations for the discrepancies between our study and that by Kamada et al. are differences in the microbiome and strains of mice used that could complicate interpretations. Future work to understand the complexities of these relationships that can yield different results are needed.

There are multiple *ler::Km* insertional mutants (sometimes confusingly annotated at Δ*ler*) that are used in the *C. rodentium* pathogenesis and immunity field (*31, 46*). The strain used in this study is from (*46*) (see methods). Our data show that at some unknown event in its lineage (before or after the generation of the *ler* mutation), this strain acquired mutations in LPS genes. LPS mutants were partially attenuated *in vivo*, though LPS-mutant challenged mice were protected from secondary challenge against wild-type *C. rodentium*. It’s likely that mutations in these LPS genes may also alter pathogen sensitivities against extracellular stress or host defenses (*47, 48*). Single mutations in *wfaP* or *rfaK* resulted in reduced antibody recognition. In other Gram-negative bacteria, spontaneous mutations in *rfaK* can arise in the population, because of antibiotic stress or bacteriophages (*49*). Consistent with is notion, we found that the LPS mutants were highly resistant to *C. rodentium* specific CrRp3 phage (*50*) (data not shown). Our work is a reminder of the importance to rigorously validate strains used in microbiological studies when making claims about gene functions including sequencing strains, complementing mutations and making clean deletions. As we do not know when in the lineage of this *ler::km* strain the mutations occurred, we encourage investigators in the *C. rodentium* field to check their strains, and if appropriate, consider reassessing some of their phenotypes and conclusions.

Kamada et al. demonstrated in C57Bl/6 mice that avirulent *C. rodentium* eventually gets excluded from the intestine because it is outcompeted by the microbiome (*51*). However in the current study, we demonstrate that phenotypically and genetically avirulent *C. rodentium* persists in the host as indicated by persistent fecal shedding. Differences in mouse stains, diets, and facilities may lead to differences in microbiome competitive effects with avirulent *C. rodentium* that can dictate whether the *C. rodentium* will persist or be outcompeted (*33, 52–55*). Indeed, in our previous work, we demonstrated in C3H/HeJ mice that dietary iron did not affect fecal shedding levels of *C. rodentium* (*20*). In the current study we show that fecal shedding in iron fed mice is reduced compared to infected mice fed control diet in C3H/Snell mice. Furthermore, the work performed for the Sanchez *et al*. study was done in a different vivarium room than the current study, which further supports that in addition to mouse strain background, different environments can influence phenotypes of *C. rodentium* mouse infections, presumably through differences in the microbiome. Future work is needed to better understand how different microbiotas and mouse strains influence aspects of *C. rodentium* infection under normal and different dietary conditions including enriched iron.

How asymptomatic carriage of a pathogen influences the ability of a host to defend against subsequent challenges with the same pathogen is not understood. In our study, we demonstrate that animals that become asymptomatically infected with virulent, are protected from subsequent virulent challenges. We demonstrated that this protection was dependent on LEE encoded virulence factors during the primary infection. We propose a model whereby systemic IgG responses that sustain asymptomatic infection with the primary infection via the selective exclusion of phenotypically virulent pathogen, also mediate protection against subsequent virulent challenges with the same pathogen, revealing a conceptually novel live vaccination strategy. Although much effort has been used to create human vaccines against EHEC and EPEC (*56*), there is still no commercially available and effective human vaccine against EPEC or EHEC. Vaccines would be a potent tool to prevent juvenile diarrheal illnesses around the world. Common vaccine strategies have been the creation of live attenuated, protein conjugates, anti-toxins, as well as “bacterial ghost” vaccines (*57, 58*). Our data suggests that long-term immunization is possible, but requires the pathogen to retain some of its virulent behavior.

Therefore, strategies which disrupt LEE PAI may be deleterious in the goal for a robust oral vaccine. We posit that a combination of mutations which sensitize A&E pathogens to the immune system plus dietary interventions to promote anti-virulence defenses to further attenuate the pathogen may provide long lasting immunity.

## ACKNOWLEDGEMENTS

This work was supported by NIH awards DPI AI144249 and R01AI14929 (JSA). This work was supported by the Hillblom Foundation Fellowship Grant (GYC). This work was supported by the NGS Core Facility of the Salk Institute with funding from NIH-NCI-CCSG: P30 014195, the Chapman Foundation and the Helmsley Charitable Trust. This work was supported by the Waitt Advanced Biophotonics core Facility of the Salk Institute with the funding from NIH-NCI: P30 014195. This work is supported by Galaxy Australia, a service provided by the Australian Biocommons and its partners. The service receives NCRIS funding through Bioplatforms Australia and the Australian Research Data Commons (https://doi.org/10.47486/PL105), as well as The University of Melbourne and Queensland Government RICF funding. We thank Nunez Lab (U Michigan) for providing the *ler::km* strain, the Sperandio lab (currently at U of Wisconin-Madison, formally at UT Southwestern) for advise on attachment assays and Geoffrey Wahl (Salk) for gifting the mCherry expression vector.

## AUTHOR CONTRIBUTIONS

GYC – Conceptualization, experimental design, data collection, data analysis, writing manuscript NRT – Data collection, data analysis

AJM – Data collection, data analysis YCL – Plasmid construction

JSA - Conceptualization, experimental design, data analysis, writing manuscript

## MATERIALS AND METHODS

### Animals

All animal experiments were done in accordance with The Salk Institute Animal Care and Use Committee and performed at Salk’s AALAC-certified vivarium. C3H/HeJ (RRID:IMSR_JAX:000659), C3H Snell (RRID:IMSR_JAX:000661), C3H SCID (RRID:IMSR_JAX:001131), C57Bl/6 (RRID:IMSR_JAX:000664), muMt^-^ (RRID:IMSR_JAX:002288) and Rag^-/-^ (RRID:IMSR_JAX:002216) were purchased from Jackson Laboratory. IgA^-/-^ mice were a generous gift from Meghan Koch (U of Wash). C3H Snell and C3H SCID mice were cross-bred to generate heterozygous and homozygous SCID mice.

### Bacteria

Wild-type *C. rodentium* DBS100 was purchased from ATCC. Bacteria were grown either on Luria Broth (LB) or MacConkey media at 37°C supplemented with antibiotics when appropriate: kanamycin (Km, 100µg/ml), chloramphenicol (Cm, 20µg/ml), or carbenicillin (Cb, 200µl/ml). The *ler::km* strain provided by the Nunez lab was from (*46*) (Personal communication with Dr. Gabriel Nunez, U Michigan). Consistent with our results reported in the current study, the Nunez lab amplified *wfaP* and *rfaK* from DNA derived from their *ler* mutant stock from (*46*) and found that the bands from the two genes were mutated in the *ler::km* mutant when compared to the wild type strain (Personal communication with Dr. Gabriel Nunez, U Michigan). The *ler::km* strain used in studies (*34, 51*) are reported to be from (*31*).

pRE112 (#43828), pFCM1 (#64948), pTNS2 (#87802), pUC18R6KT-mini-Tn7T-Km (#64969) and pUC18R6K-mini-Tn7T (#64958) plasmids were purchased from Addgene. The Tn7 vector (pSAC1) with Cm resistance was generated by cloning the Cm^R^ gene from pFCM1 into pUC18R6K-mini-Tn7T by standard Gibson reaction. For cis expression in *C. rodentium*, genes were amplified by PCR and cloned into pCRII-TOPO2 behind a lac promoter. To transpose Tn7-Cm constructs into *C. rodentium*, Tn7 vectors and pTNS2 helper plasmid were co-electroporated into bacteria and selected on LB supplemented with Cm. The transduction of Tn7 transposons into the glmS site was verified by PCR. To construct in-frame deletions in *C. rodentium*, approximately 1kb regions flanking target genes were amplified and joined by Gibson reaction into pRE112. Gene deletion constructs were then electroporated into *C. rodentium* and selected by Cm. Isolates were cultured without Cm and selected on LB sucrose plates to isolate excision of the plasmid. Isolates were screened by PCR for the deleted allele. Bacterial strains and primers are listed in **Supplemental Table 2**.

### Infections

Six- to ten-week-old male mice were used for all infection experiments. Mice were purchased from Jackson labs and housed in our mouse facility for 3 days to acclimate prior to experimentation. During the acclimation period, mice were fed ad libitum standard mouse vivarium chow (normal diet for this study). For infections, *C. rodentium* cultures were grown from individual colonies on LB plates, and shaken overnight at 37°C. Cultures were then centrifuged and the pellet was resuspended in PBS. Overnight fasted mice were orally gavaged with 7.5 × 10^8^ CFUs dose of *C*. rodentium and given control or 2% carbonyl iron chow (Envigo, TD.99398 and TD.08714). Weight and fecal shedding were monitored daily. For survival, mice were evaluated clinically daily as described below and assigned a clinical score. We employed a clinical scoring system to determine when mice become moribund. As animals reach stage one they are euthanized immediately to prevent unnecessary pain and suffering. Our scoring system is outlined below: 5=Normal exploratory behavior, rearing on hind limbs, and grooming. 4=Mild: Reduced exploratory behavior, rearing on hind limbs, and grooming. Slower and/or less steady gait, but free ambulation throughout the cage. 3=Moderate. Limited voluntary movement. Slow, unsteady gait for >5 seconds. 2=Severe. No voluntary movement, but mouse can generate slow, unsteady, gait for >5 seconds. 1=Moribund. Mouse does not move away from stimulation by researcher and no longer rights itself. As mice reached stage one they were euthanized and counted as a death event. Survival curves include both mice that died as well as those that were euthanized due to reaching clinical endpoint. For protection experiments, mice were infected with *C. rodentium*. Initially mice were given a 2% carbonyl iron diet for two weeks before swapping back to normal chow. After 28 days, mice were rechallenged with Cm^R^ wildtype *C. rodentium* and kept on their normal chow ad libitum.

### Fecal Shedding

Fecal pellets were homogenized in 1mL of PBS and serially titrated across 10^0^ to 10^−5^ dilutions. Dilutions were then plated on MacConkey plates with antibiotics when necessary. Plates were incubated overnight at 37°C and colonies were quantified.

### Bacterial ELISAs

C3H SCID mice were infected with *C. rodentium* and fed 2% carbonyl iron diet. At two weeks post infection, small intestine, cecum, and colon were flushed with PBS with protease inhibitor (Sigma, P2714) and EDTA. Solids were pelleted and supernatants were frozen at -20°C until analysis. Additionally, serum was stored at - 80°C.

To perform bacterial ELISAs as described in (*34*), wild-type or mutant strains of *C. rodentium* strains were grown overnight in DMEM at 37°C. Bacteria cultures were washed once in PBS and heat-killed (HK) in a 70°C water bath for 1 hour. HK-bacteria were incubated in 96-well ELISA plates overnight at 4°C. Plates were washed with PBS+BSA, 1:10 lumenal or 1:1000 serum primary antibody, then 1:1000 anti-IgG/A/M conjugated to horseradish peroxidase. Plates were thoroughly washed with PBS with tween-20. Antibody binding was visualized by o-phenylenediamine dihydrochloride reaction at 492nm absorbance on a microplate reader.

### Attachment assay

The attachment assay, aka fluorescent actin staining (FAS) assay, was performed according to (*59*)mCherry labeled bacteria were grown overnight in LB. HeLa cells were grown in 24-well plates with round coverslips at a density of 2×10^5^ cells per well. Prior to infection, both bacteria and cells were washed with PBS and 50μl of bacteria were added per well. Infected cells were incubated at 37°C and 5% CO2 for 4 hours washed once with PBS and HeLa cell media. Infected cells were incubated for an additional 4 hours before fixation with 4% PFA, permeabilization with triton-X100, and actin staining with phalloidin-alexa488. Bacterial attachment was visualized on a LSM880 with Airyscan. Airyscan, stitching and maximum intensity projection processing were performed using the Zeiss Zen software (https://www.zeiss.com/microscopy/en/products/software/zeiss-zen.html). Micrograph color contrast and image cropping were performed using FIJI (*60*) (https://imagej.net/software/fiji/).

### LPS analysis

*C. rodentium* LPS was extracted by hot-phenol as outlined in (*61*). LPS extracts were then run in a 12% polyacrylamide gel and stained with ProQ Emerald 300 LPS gel stain kit (Thermo, P20495). Stained gels were visualized on a UV transluminator and a Nikon D3000 dSLR camera equipped with a Kodak yellow #15 filter.

### Genome sequencing/analysis

Bacteria were grown overnight in LB cultures and then genomic DNA was extracted using standard phenol chloroform protocol. Genomic samples were checked for quality by nanodrop, Qbit and bioanalyzer. Both Illumina and Oxford nanopore libraries were constructed by the Salk NGS core and run on either and Illumina MiSeq (2×150 paired-end) or Oxford Nanopore GridION x5, respectively. Galaxy Australia suite of tools for genome analysis (https://usegalaxy.org.au/). Unicycler (*62*) was used for de novo hybrid genome assembly.

Genome alignments were analyzed using the Artemis Comparison Toll (ACT) (*63*). To identify novel insertion sequence (IS) events in *C. rodentium*, ISFinder (https://isfinder.biotoul.fr/) BLAST tool was used to identify native *C. rodentium* IS. IS queries were assessed by running Illumina reads from different strains through ISMapper (*64*)(https://github.com/jhawkey/IS_mapper).

### qPCR

To measure bacterial gene expression, bacteria were first grown to exponential phase in LB or LB+ lactose cultures. Cells were pelleted then immediately frozen in liquid nitrogen. Nucleic acids were extracted using TRIzol™ (Thermo, 10296028), mechanical bead beating, and standard ethanol precipitation. Following nucleic acid extraction, RNA was purified by DNase treatment, then cDNA was produced using the Superscript™ IV Reverse Transcriptase kit (Invitrogen, 18090050). Gene expression was measured using iTaq™ Universal SYBR^®^ Green Supermix (Biorad, 1725125) in an Applied Biosystems QuantStudio 5 Real-Time PCR machine. qPCR primers and housekeeping gene (*65*) are listed in **Supplemental Table 2**.

### Graphics and statistics

Statistical analysis and graphing were performed using Graphpad Prism 9. Normality tests were run to determine the distribution of the data and unpaired t-test, Mann-Whitney test, One Way ANOVA with post Tukey test, Kruskal Wallis with post Dunn’s test or Two Way ANOVA were performed for pairwise comparisons and as indicated in the figure legends. Log rank analysis for survival. Sample sizes and information about replicates are included in the figure legends. Inkscape (https://inkscape.org/) and Biorender (https://biorender.com/) were used for producing graphics describing experimental overview and model. Open-source icons were obtained from flaticon.com and reactome.org.

### Data deposit

The data have been deposited with links to BioProject accession number PRJNA926120 in the NCBI BioProject database (https://www.ncbi.nlm.nih.gov/bioproject/).

## SUPPLEMENTAL FIGURE LEGENDS

**Supplemental Figure 1.**
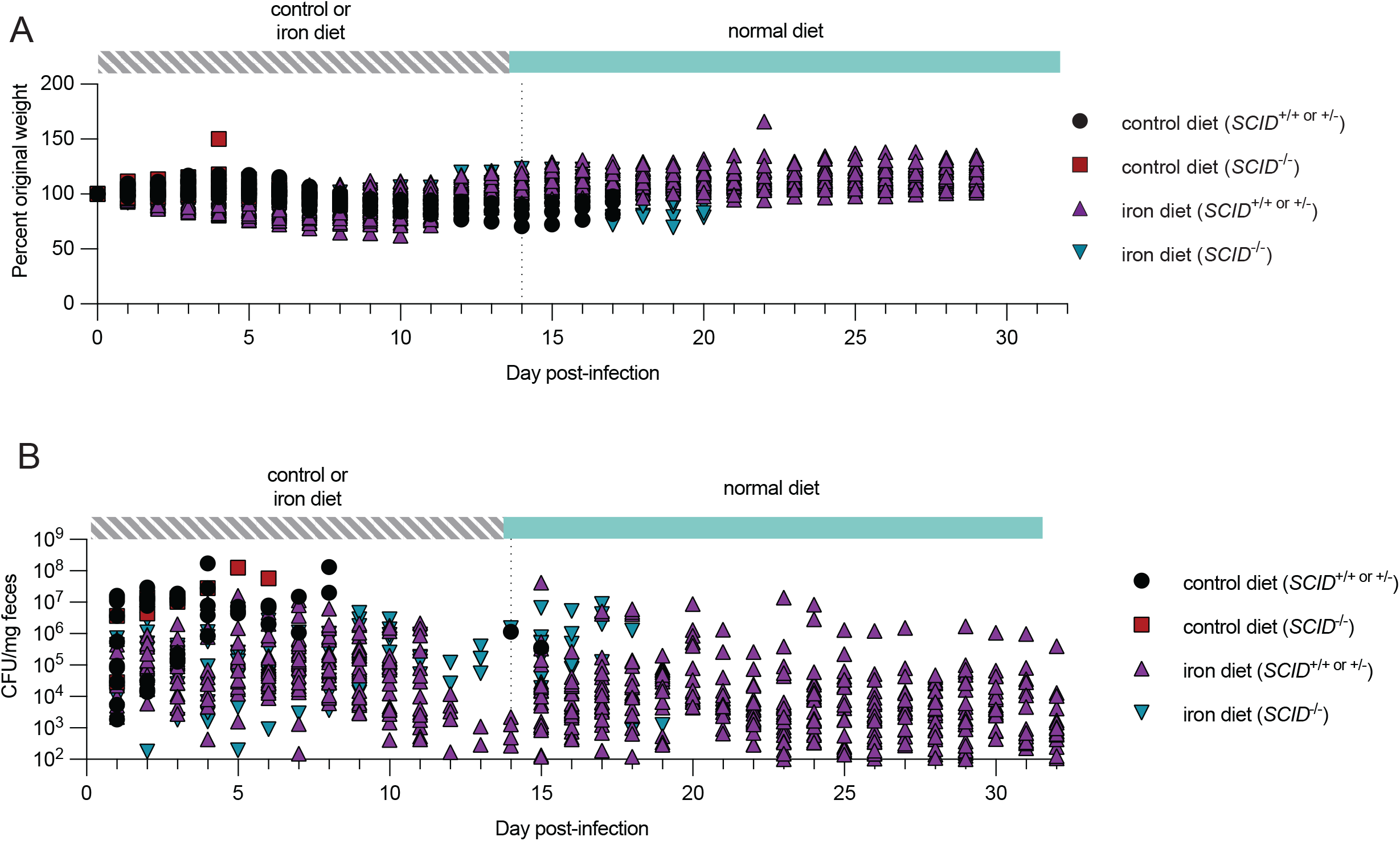
Adaptive immunity is required for asymptomatic carriage of *C. rodentium*. (A) Percent original weight data from Figure 1C and (B) fecal shedding data in Figure 1C plotted as individual data points.

**Supplemental Figure 2.**
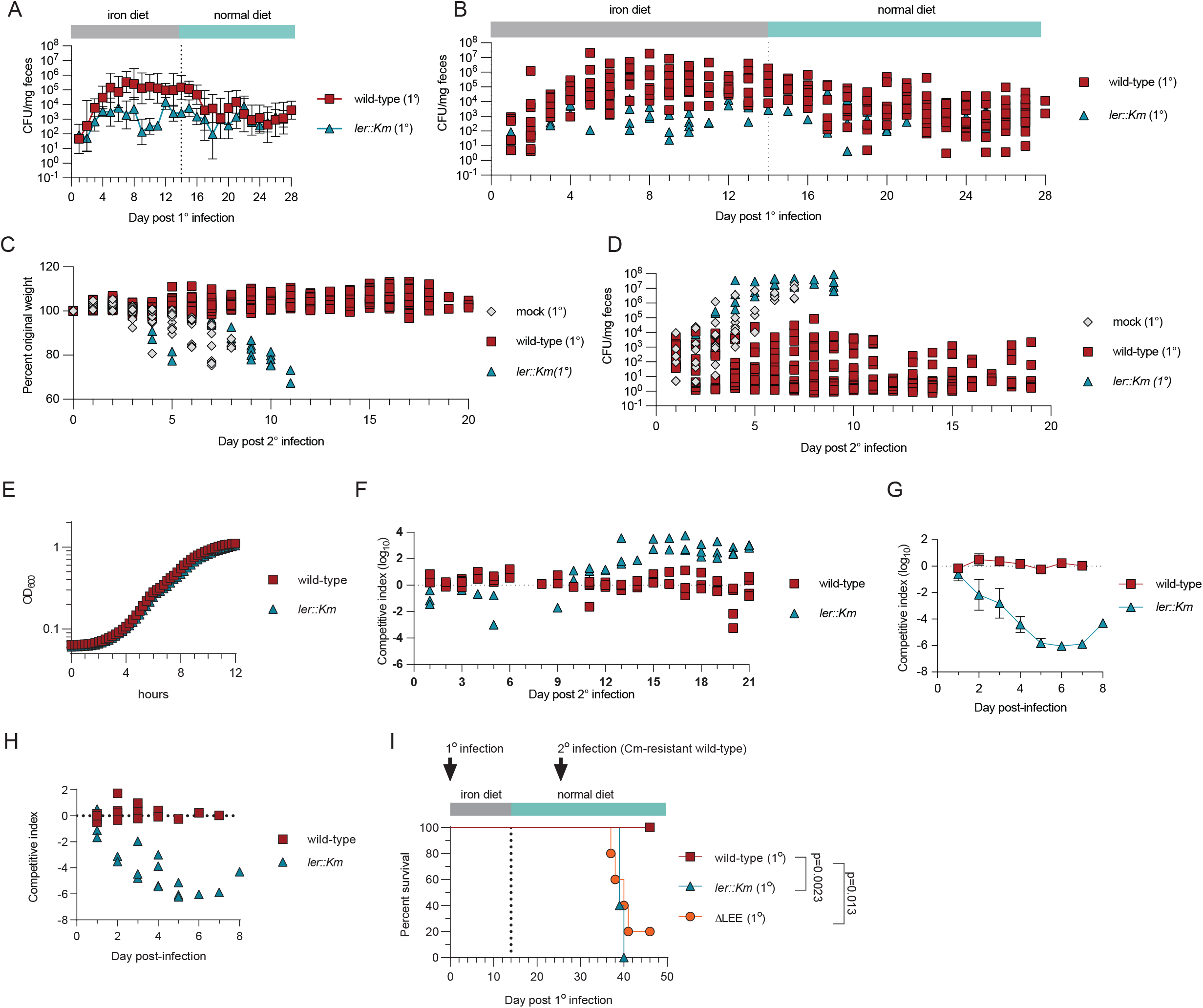
Asymptomatic carriers are protected from subsequent challenges with *C. rodentium*. (A) Fecal shedding average or (B) individually plotted data points of primary infection strain, wild-type or *ler::Km* from experiment in Figure 2B. (C-D) Individually plotted data points from Figure 2C-D, respectively. (E) Growth curve of wild-type or *ler::Km* in LB media. n = 4 per condition (F) Individually plotted data points from Figure 2F. (G) The average or (H) individually plotted data points of competition experiment in C3H/HeJ mice challenged with 1:1 CFU of wild-type or *ler::Km* over wild-type. n = 5 mice per condition. Data represent one biological replicate. (I) Protection experiment, as outlined in Figure 2A, but including ∆LEE strain during primary infection. n = 5 mice per condition. Data represent one biological replicate. Geometric mean +/-geometric SD for (A), +/-SEM for (G). Log rank analysis for (I).

**Supplemental Figure 3.**
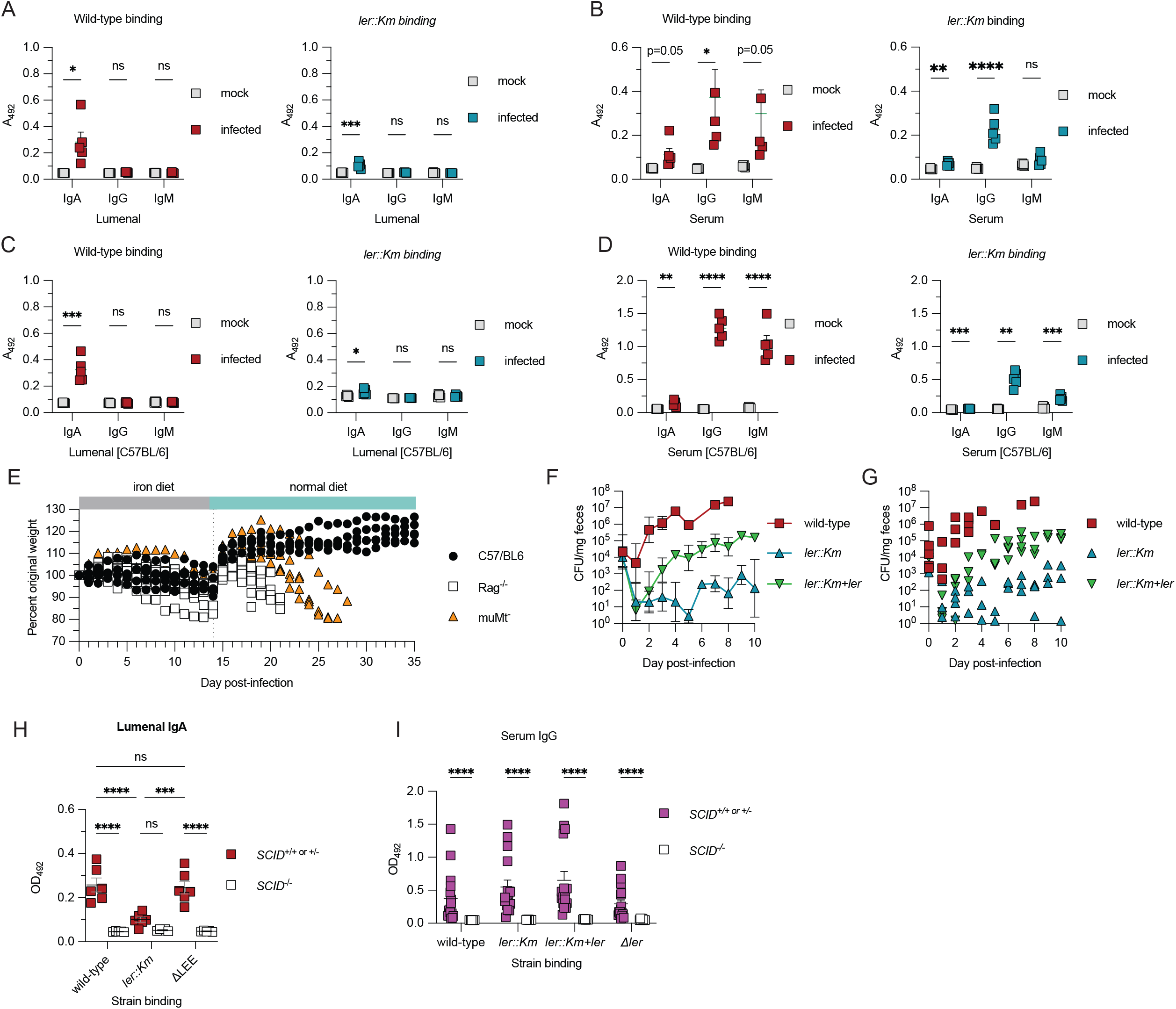
*C. rodentium* specific antibodies do not distinguish between wild-type and Ler/LEE deficient strains. Whole bacteria ELISAs quantifying (A&C) lumenal or (B&D) serum IgA, IgG or IgM antibodies binding to wild-type or *ler::Km C. rodentium*. Samples were collected from C3H mice (A & B) or B6 mice (C & D) mock (PBS) or wild-type *C. rodentium* infected mice fed iron diet at two weeks post infection n = 5 mice per condition. Data represent one biological replicate. (E) individually plotted data points of Figure 3D. (F) average or (G) individual data point fecal shedding of C3H mice infected with wild-type, *ler::Km*, or *ler::Km+ler*. n = 4-5 mice per condition. Data represent one biological replicate. Whole bacterial ELISAs quantifying (H) lumenal IgA or (I) serum IgG binding against wild-type, *ler* mutant, or ∆LEE stains. n = 5 to 16 samples per condition. Lumenal and serum samples were collected from wild-type *C. rodentium* infected C3H SCID mice fed iron diet for two weeks post-infection. Data represent one biological replicate. Mean +/-SEM or Geometric mean +/-geometric SD. Unpaired t-test, Mann Whitney test or One Way ANOVA with post-Tukey test for pairwise comparisons

**Supplemental Figure 4.**
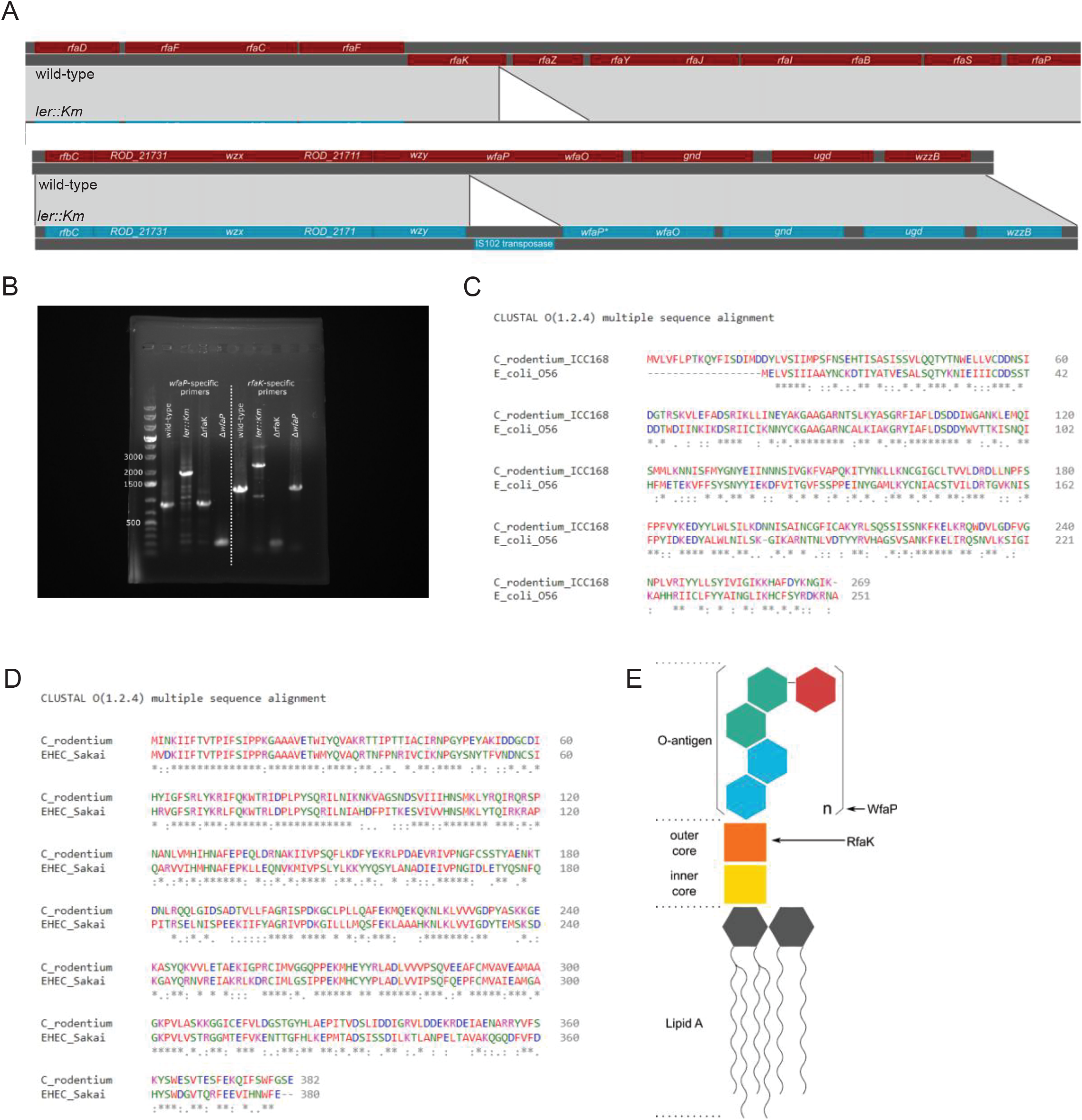
LPS, not ler, is recognized by *C. rodentium* specific antibodies. (A) Genome alignments of the *rfa* and *wfa* operons from wild-type or *ler::Km* genome assemblies using the Artemis Comparison Tool (ACT). (B) PCR reactions of wild-type, *ler::Km*, and LPS mutants amplifying *wfaP* or *rfaK* genes. (C-D) Protein sequence alignments of *C. rodentium* (C) *wfaP* and (D) *rfaK* against *E. coli O56 wfaP* and *EHEC Sakai rfaK*, respectively. (E) Diagram of WfaP and RfaK enzymatic activity during LPS biosynthesis.

**Supplemental Figure 5.**
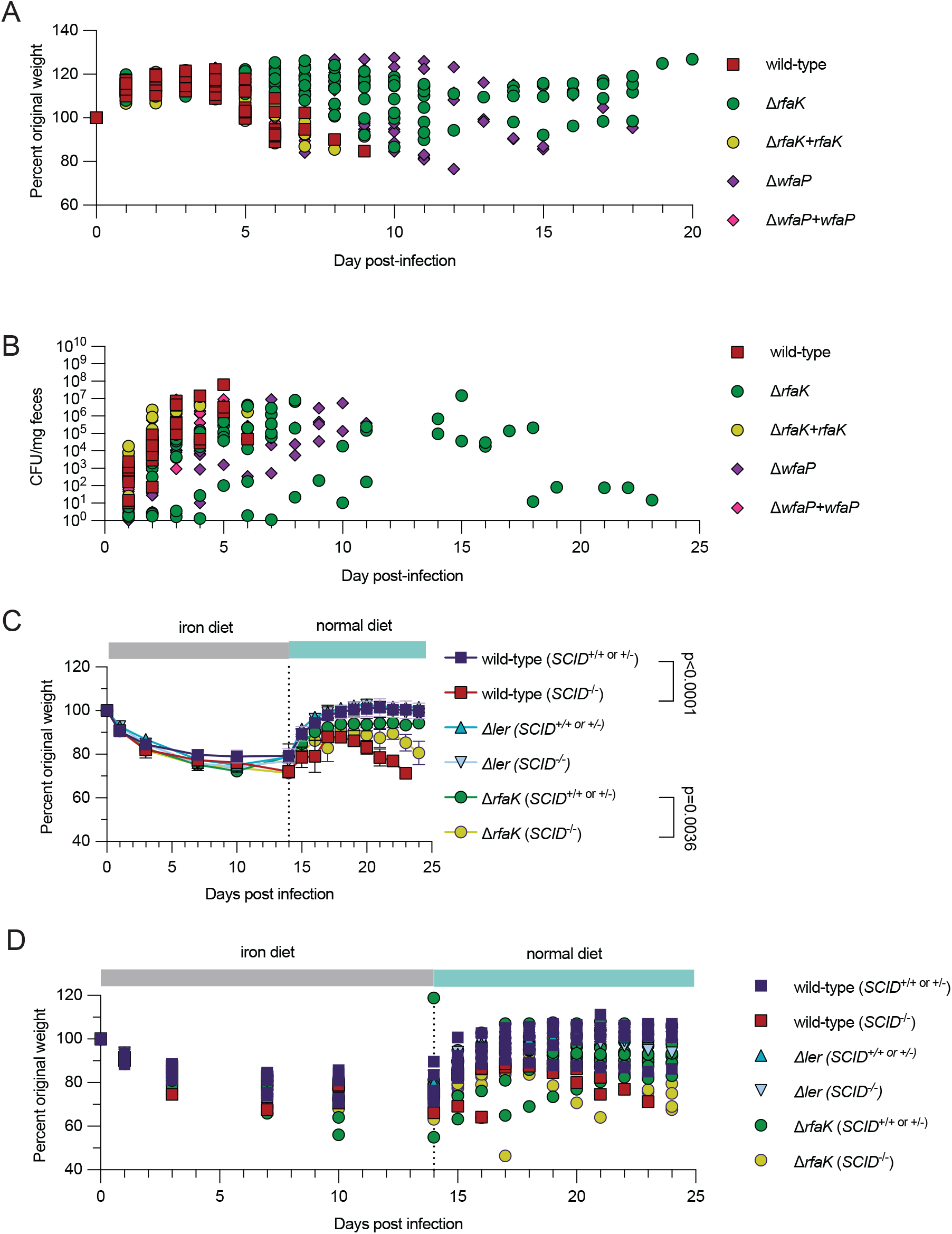
LPS promotes virulence of *C. rodentium*. (A) Percent original weight and (B) fecal shedding of *C. rodentium* infected mice as shown in Figure 5C and D but plotted individual data points. (C-D) Percent original weight as (C) average and (D) individual data points of mice in Figure 5E. Wild-type (*SCID*^*+/+ or +/*-^) = 5 mice, wild-type (*SCID*^*-/-*^) = 3 mice, *Δler* (*SCID*^*+/+ or +/-*^) = 2 mice, *Δler* (*SCID*^*-/-*^) = 3 mice, Δ*rfaK* (*SCID*^*+/+ or +/-*^)= 9 mice, Δ*rfaK* (*SCID*^*-/-*^) = 7 mice. Data represent two biological replicates combined. Error bars indicate +/-SEM for weight curve.

**Supplemental Figure 6.**
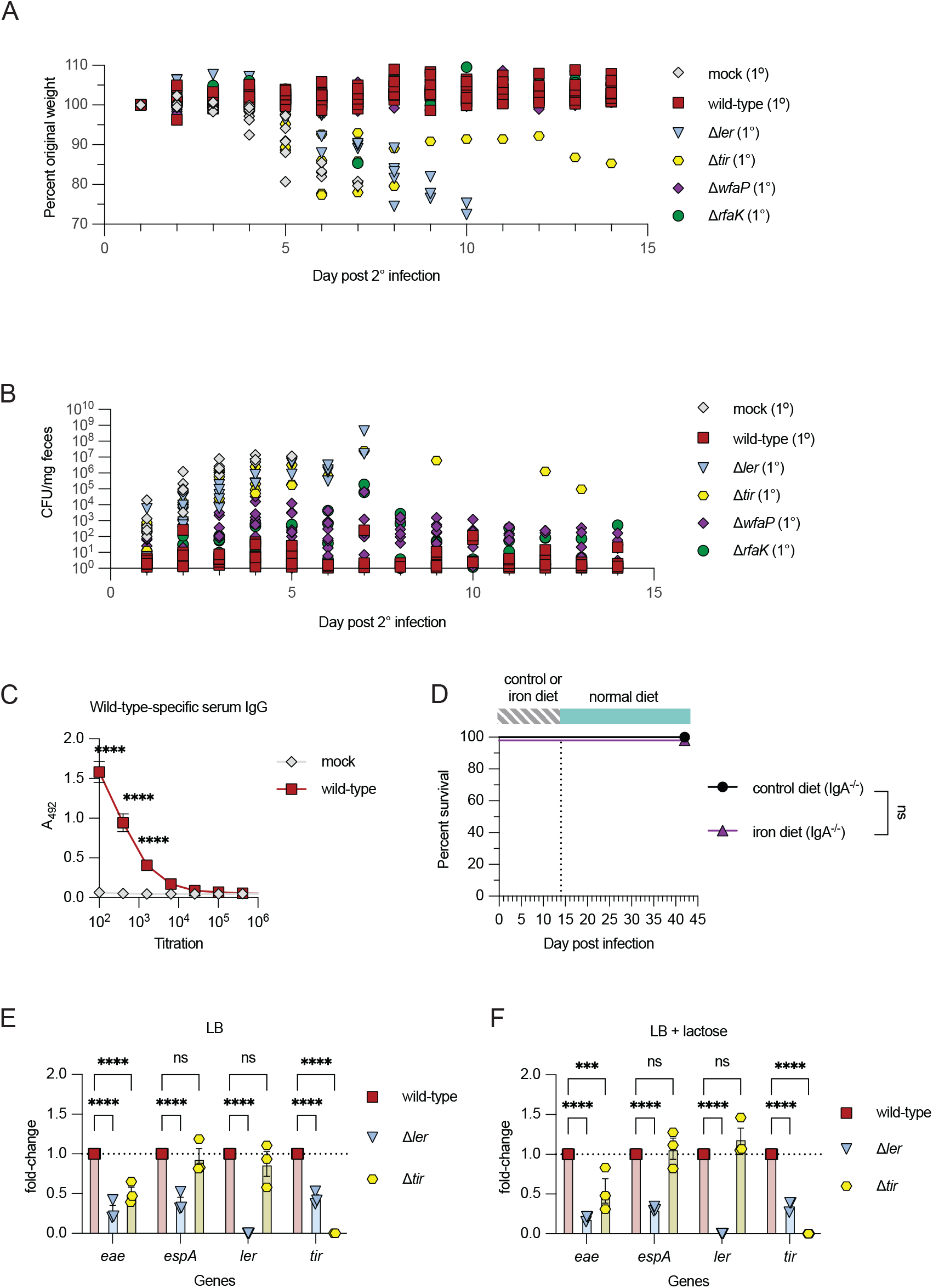
Pathogen behavior is necessary for protection of asymptomatic carriers with subsequent infections. (A) Percent weight loss and (B) fecal shedding of *C. rodentium* infected mice as shown in Figure 6D & E but plotted individual data points. (C) Whole bacteria ELISA examining titrated serum IgG, collected from wild-type or mock-infected mice, against wild-type *C. rodentium*. n = 10 mice per condition. Data represent one biological replicate. (D) *IgA*^*-/-*^ mice were infected with wild-type *C. rodentium* and given control or iron chow for two weeks and then given their normal chow. Survival was monitored. n = 5 mice per condition. Data represent two biological replicates combined. (E-F) qPCR analysis of virulence genes *eae, espA, ler*, and *tir* expressed in wild-type, ∆*ler*, and *∆tir* grown in (E) LB or (F) LB supplemented with 0.2% lactose. n = 3 mice per condition. Data represent three biological replicates combined. Statistical significance was calculated using two-way ANOVA for unpaired t-test or pairwise comparisons and Log rank analysis for survival. Error bars +/-SEM. \*\*\**p* < 0.001, \*\*\*\**p* < 0.0001.

**Supplemental Table 1.**
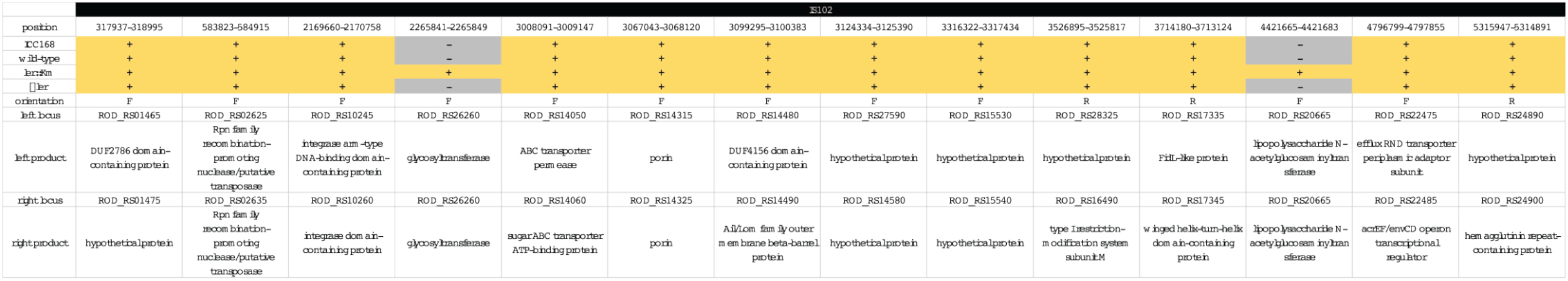
ISMapper analysis of IS102 transposon insertions of sequenced strains.

**Supplemental Table 2.**

Bacterial strains, plasmids and primers

